# Modified split application of nitrogen with biochar improved grain yield and nitrogen use efficiency in rainfed maize grown in Vertisols of India

**DOI:** 10.1101/2020.07.13.200345

**Authors:** Bharat Prakash Meena, Pramod Jha, K. Ramesh, A.K. Biswas, R. Elanchezhian, S. Neenu, H. Das, A.O. Shirale, Ashok K. Patra

## Abstract

Conventionally, non-judicious and blanket fertilizer nitrogen (N) used in rainfed maize lead to higher N losses, low N use efficiency (NUEs) and poor yields due to substandard agronomic management practices. To avoid such N losses, fertilizer additions are synchronized with plant uptake requirements. In this context, agronomic based management focused on optimizing N rates and biochar application is essential for improved NUEs and crop productivity. Keeping this in view, a field experiment was conducted during 2014, 2015 and 2016 in rainfed maize (*Zea mays* L.) grown in Vertisols of India. In this study, twelve treatments that comprised of N omission plot (N0), skipping of basal rate, multi-split topdressing at varying time as broadcast and band placement, soil test crop response (STCR) based NPK with target yield 6.0 t ha^-1^ in maize and biochar application (10 t ha^−1^) were investigated. The experiment was conducted following a Randomized Complete Block Design (RCBD) set up with three replications. Pooled analysis of three years data revealed that the application of N rates (120 kg Nha^−1^) in 2 equal splits (60 kg Nha^−1^) at knee high (V8) and tasseling (VT) stages with skipped basal N rate, achieved higher maize grain yield (5.29 t ha^−1^) ascribed to the greater growth parameters, yield components and N uptake compared to the recommended practices. Biochar application (10 t ha^−1^) as soil amendments along with multi top dressed N (120 kg N ha^−1^) into 3 splits also increased the grain yield. Delayed N application at V8 and VT growth stages, resulted in higher N uptake, agronomy efficiency (AE), partial factor productivity (PFP), physiology efficiency (PE) and recovery efficiency (RE). Biochar along with N fertilizer also improved the soil organic carbon (5.47g kg^−1^), ammonium-N (2.40 mg kg^−1^) and nitrate-N (0.52 mg kg^−1^) concentration in soil (P<0.05) as compared to non-biochar treatments. Application of biochar along with chemical fertilizer (120 kg Nha^−1^) significantly increased the concentration of ammonium (2.40 mg kg^−1^) and nitrate (0.52 mg kg^−1^) in soil (P<0.05) as compared to non-biochar treatments. The perfect positive linear relationship illustrated that the grain yield of rainfed maize was highly dependent (*R*^*2*^=0.99 at p<0.0001) on N availability, as indicated by the fitted regression line of maize grain yield on N uptake. On the other hand, factor analysis revealed, the one to one positive function relationship of biomass with N uptake at V8 and VT growth stages. Principal Component Regression (PCR) analysis showed that PC1 acted as a major predictor variable for total dry matter yield (TDMY) and dominated by LAI and N uptake. Consequently, these results expressed that the agronomic management based multi-top dressed N application and biochar application to achieve higher yield and greater NUEs in rainfed maize is strongly linked with N application into splits.

## Introduction

Globally maize (*Zea mays* L.) is an important food crop with highest productivity compared to any other food crops [1]. Since, maize is a heavy nutrient exhaustive crop [2] and being C4 plant type, requires a regulated and assured supply of nutrient particularly nitrogen (N) fertilizer throughout its growing period [3]. Nitrogen fertilization assured centerstage for maize production [4] but N fertilizer is a costly input and it is very complicated to manage because its utilization is largely dependent on agronomic, genetic, biological, soil and climate factors [5]. Nitrogen fertilization and time of its application is most crucial for higher productivity of maize [6]. However, non-judicious and excess use of chemical N fertilizer to cropland may have negative impact on air, water, soil and biodiversity and also generate green house gases [7]. Besides, conventional practice of N application (large portion) through surface broadcasting just after maize planting instead of split application have led to the decrease in the crop production and low N use efficiency in India [8] because the N is applied to the soil is vulnerable to losses from the soil– plant system passing through volatilization, de-nitrification, leaching, run-off causing serious threat to environmental quality [9]. Thus, full supply/application of N at the time of planting has resulted in poor matching between the soil supply and the crop demand, leading to high organic N in soil before the peak crop requirement [10], resulting into unpromisingly low NUE. The poor NUE not only have negative impact on the crop yield but also have raised the environmental concerns [11]. Therefore, efficient N fertilizer management strategies will be at the forefront of measures to improve crop yield and NUE [12]. Numerous studies have pioneered on improved N management practices to achieve higher crop yield and NUE in maize such as split application [13], timing N of application [14], band placement of N fertilizer [15], site specific nitrogen management [16] and soil test based N rate [17]). These agronomic management practices either boost the availability total amount of N, tuned as a multi-split application through better synchronization or decrease the N losses ([18].Top dressing N into multi-split application increased total N accumulation and decreased stem nutrient content remobilization [19]. Even though, uncontrollable factors like temperature, rainfall timing, intensity and amount, and interactions of temperature and rainfall were also strong relationship with crop yield and N uptake [20]. Skipping the basal rate of N and sifting it at crop establishment stage enhanced the grain yield and agronomic use efficiency at 130 kg N ha^−1^ application in rice [21]. Maize produced higher growth and yield component with multi-split N application at later growth stage and also had a positive impact on maize yields [22]. With the application of N at V6 stage, plant exhibited N deficiency resulting in decreased cell division and cell elongation, which decreased leaf length and delayed time for leaf expansion and consequently decreased grain yield [23]. In general, it was reported that the benefit of late split N applications is mainly depended on quantity and application pattern and there were no evidence of decline in yield level with delayed N application in maize until V11 [24].

Most of previous studies on multi-split N application for improving crop yield and N use efficiencies in rainfed maize are limited and also N recommendations for maize are less accurate than desired. The objectives of this study were to; (i) determine the effect of multi-split N application on biomass and N accumulation at different growth stages, (ii) know the time and rate of N application for better synchrony between soil N supply and crop N demand for potential yield and greater NUEs, (iii) identify the impact of skipping basal N rate on plant growth and N uptake in whole plant biomass, (iv) investigate the impact of biochar application as a soil amendment on crop yield and N use efficiencies in rainfed maize.

## Materials and methods

### Description of experimental site

A 3–years field study was performed in 2014, 2015 and 2016 at the research farm of ICAR-Indian Institute of Soil Science (ICAR–IISS), Bhopal, Madhya Pradesh, India (23°18′N, 77°24′ E, 485 m above mean sea level). The soil of the experimental site was an Isohyperthermic, Typic Haplustert with deep heavy clayey in texture (24.5% sand, 23.5% silt and 52.0% clay) and slightly alkaline in reaction (pH=7.8). The soil was low in organic carbon (4.5 g kg^−1^), low in available KMnO4–N (206.6 kg ha^-1^), but high in available Olsen’ P (50.0 kg ha^-1^) and high in NH4OAc–K status (621.0 kg ha^-1^) (Table 1). The climate of the region is sub-humid with cool, dry winters, a hot summer and a humid monsoon season with 30 years mean annual rainfall of 1130 mm and potential evapo-transpiration of 1400 mm [25].

**Table 1.**
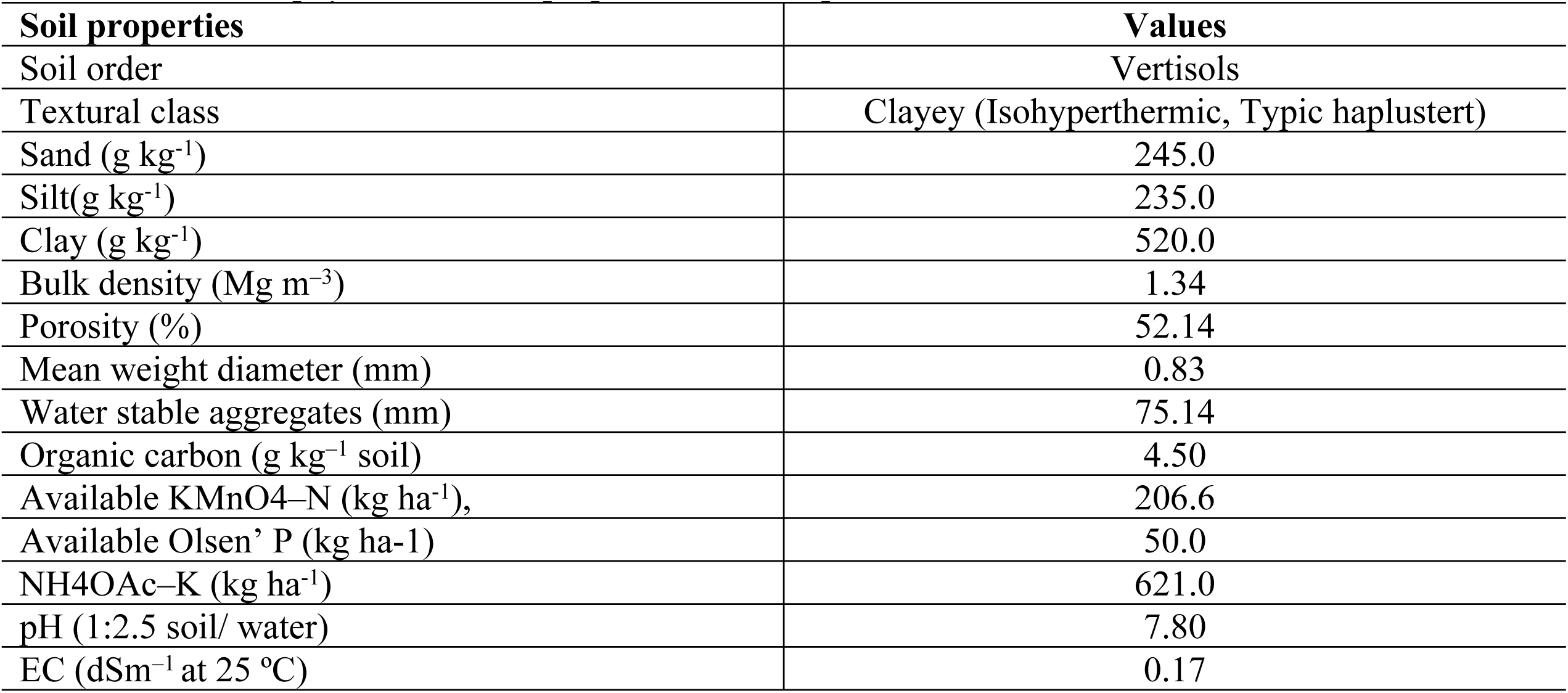
Initial soil physicochemical properties of the experimental site

| Soil properties | Values |
| --- | --- |
| Soil order | Vertisols |
| Textural class | Clayey (Isohyperthermic, Typic haplustert) |
| Sand (g kg<sup>-1</sup>) | 245.0 |
| Silt(g kg<sup>-1</sup>) | 235.0 |
| Clay (g kg<sup>-1</sup>) | 520.0 |
| Bulk density (Mg m<sup>-3</sup>) | 1.34 |
| Porosity (%) | 52.14 |
| Mean weight diameter (mm) | 0.83 |
| Water stable aggregates (mm) | 75.14 |
| Organic carbon (g kg<sup>-1</sup> soil) | 4.50 |
| Available KMnO<sub>4</sub>-N (kg ha<sup>-1</sup>), | 206.6 |
| Available Olsen' P (kg ha<sup>-1</sup>) | 50.0 |
| NH<sub>4</sub>OAc-K (kg ha<sup>-1</sup>) | 621.0 |
| pH (1:2.5 soil/ water) | 7.80 |
| EC (dSm<sup>-1</sup> at 25 °C) | 0.17 |

Daily fluctuation in rainfall, minimum (Tmin) and maximum temperature (Tmax) during the study period are presented in Figs 1–3. The maize crop was grown during wet season (rainy) from June to October. The weather parameters during experimentation were recorded at ICAR-IISS metrological observatory near to the experimental plot. The meteorological data showed a marked variation in weather condition during the three years of the experiment. Temperatures (maximum and minimum) remained more or less constant while, rainfall varied significantly during crop season.

**Fig. 1.**
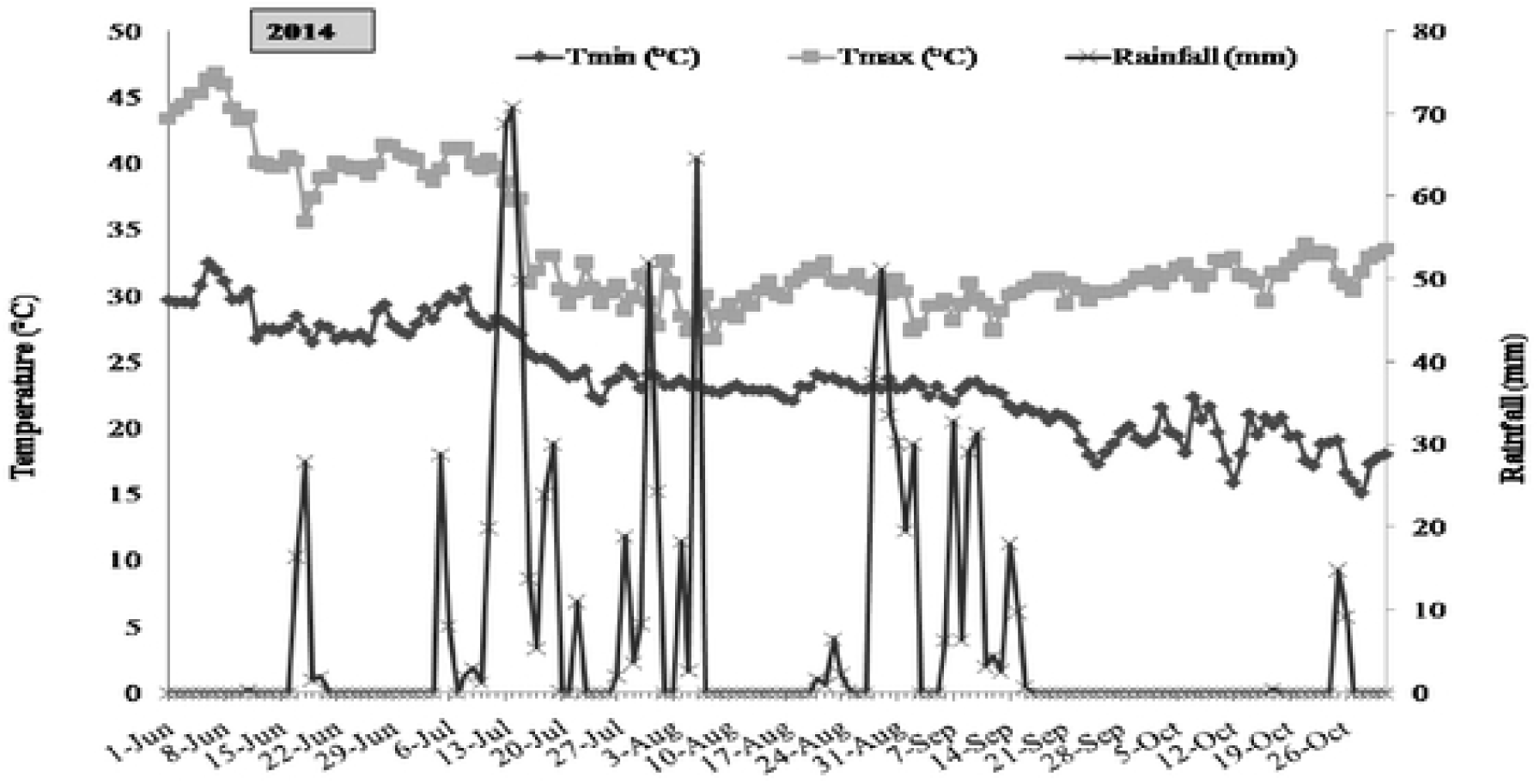
Daily fluctuation in rainfall and temperature during the crop season of 2014.

**Fig. 2.**
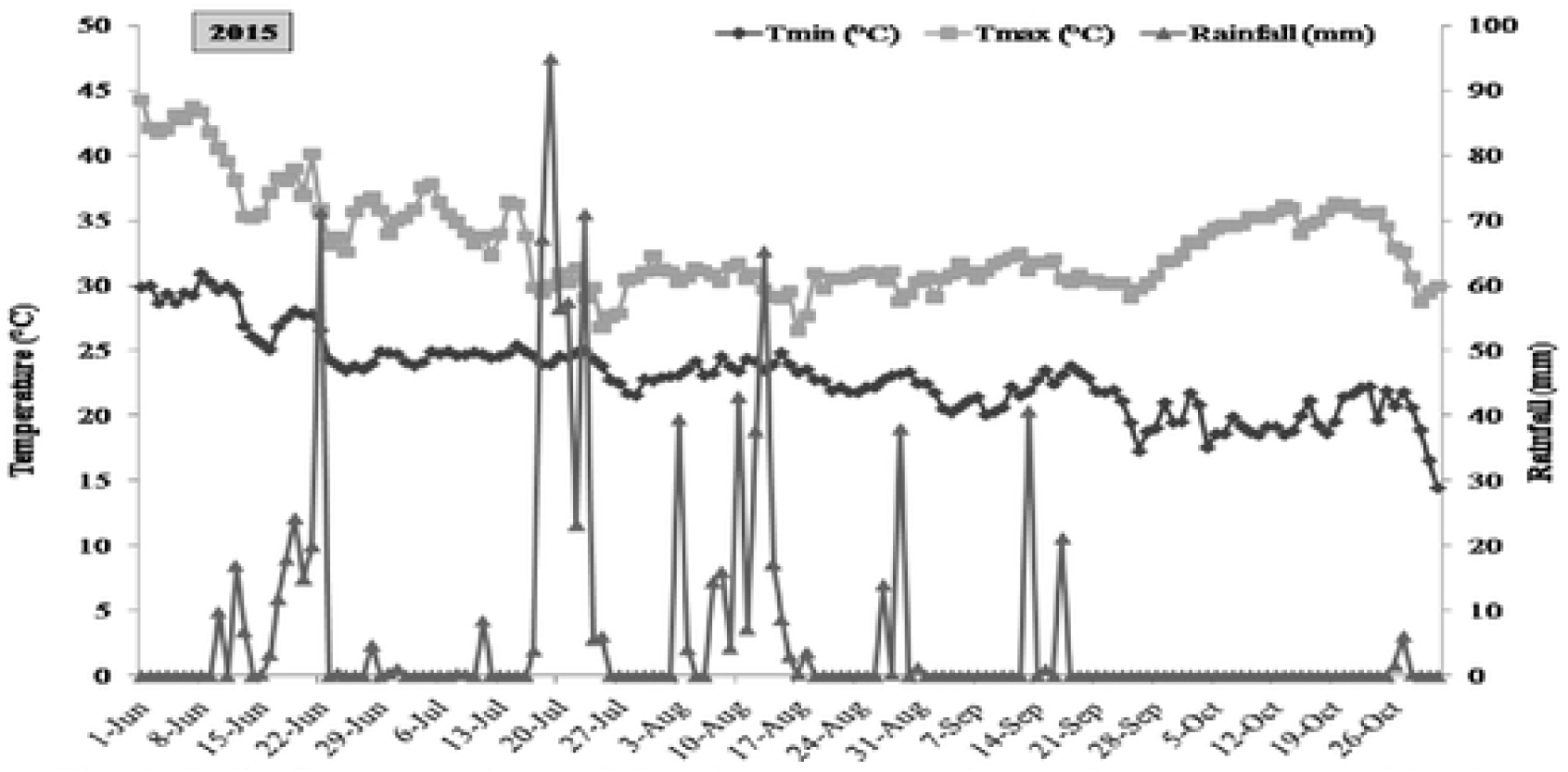
Daily fluctuation in rainfall and temperature during the crop season of 2015.

**Fig. 3.**
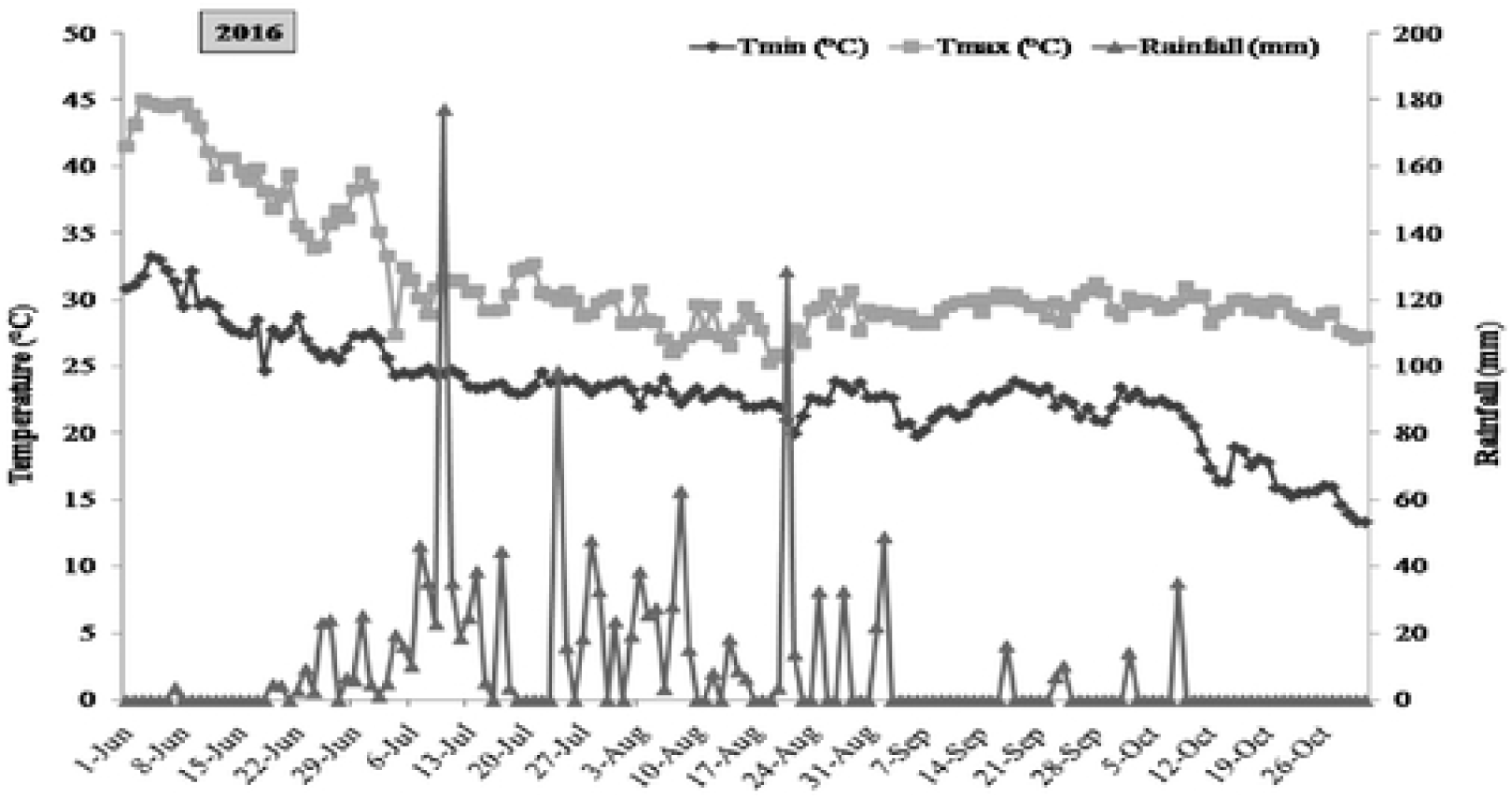
Daily fluctuation in rainfall and temperature during the crop season of 2016.

### Experimental set-up

The experiment was arranged with twelve treatment combinations in Randomized Complete Block Design (RCBD) for rainfed maize with 6 m x 5m plots replicated thrice during all three years. The N rates including 0, 90, 120 and 175 kg ha^-1^ (STCR) were applied at basal (at planting), knee high (V8), tasseling (VT) and silking stages (R1) and basal N rate was skipped in some treatment and it was applied at V8 and VT stages. An absolute NPK control and N control plots were also maintained. Fertilizer N rates and time of application are further described in Table 2. The N rate was applied through urea as per treatment details and all plots received the same treatments throughout the period of study. The sources of N, P, and K were prilled urea, single super phosphate (SSP) and muriate of potash (MOP), respectively. While, recommended dose of P (40 kgha^-1^), K (60 kgha^-1^) and Zn (5 kgha^-1^) were applied uniformly in all treatments except absolute control (N0P0K0) at the time of final land preparation. One treatment was kept on soil test crop response (STCR) equation based NPK fertilizers 175:57.6:55.4 kg ha^−1^N, P_2_O_5_ and K_2_O, respectively for a target yield of 6.0 t ha^−1^.

**Table. 2.** Treatment description and fertilization regime for the maize crop used in all three years

| Treatments |  | N rate (kg ha<sup>-1</sup>) at different growth stages |  |  |  | Total Nitrogen (kg ha<sup>-1</sup>) |
| --- | --- | --- | --- | --- | --- | --- |
|  |  | Basal | Knee high (V8) | Tasseling (VT) | Silking (R1) |  |
| T1 | Absolute control | 0 | 0 | 0 | 0 | 0 |
| T2 | N control (N<sub>0</sub>) | 0 | 0 | 0 | 0 | 0 |
| T3 | N<sub>60</sub>-N<sub>30</sub>-N<sub>30</sub> through Broadcast | 60 | 30 | 30 | 0 | 120 |
| T4 | N<sub>30</sub>-N<sub>30</sub>-N<sub>30</sub> through Band placement (Basal) | 30 | 30 | 30 | 0 | 90 |
| T5 | N<sub>0</sub>-N<sub>60</sub>-N<sub>60</sub> through Broadcast | 0 | 60 | 60 | 0 | 120 |
| T6 | N<sub>0</sub>-N<sub>45</sub>-N<sub>45</sub> through Broadcast | 0 | 45 | 45 | 0 | 90 |
| T7 | N<sub>0</sub>-N<sub>40</sub>-N<sub>40</sub>-N<sub>40</sub> through Broadcast | 0 | 40 | 40 | 40 | 120 |
| T8 | N<sub>0</sub>-N<sub>30</sub>-N<sub>30</sub>-N<sub>30</sub> through Broadcast | 0 | 30 | 30 | 30 | 90 |
| T9 | N<sub>0</sub>-N<sub>60</sub>-N<sub>30</sub>-N<sub>0</sub> through Broadcast | 0 | 60 | 30 | 0 | 90 |
| T10 | *STCR based NPK through Broadcast (Target yield -6.0 Mg ha<sup>-1</sup>) | 86.3 | 43.2 | 43.2 | 0 | 172.7 |
| T11 | Biochar (10 t ha<sup>-1</sup>) + T3 | 60 | 30 | 30 | 0 | 120 |
| T12 | Biochar (10 t ha<sup>-1</sup>) + T4 | 30 | 30 | 30 | 0 | 90 |

\*STCR, Soil test crop response equation

Soil test crop response equation (STCR) based fertilizer rates was calculated based on initial soil test value of N, P and K with target yield and adjustment by STCR prescription equations. The fertilizer prescription equations for maize were as follows;

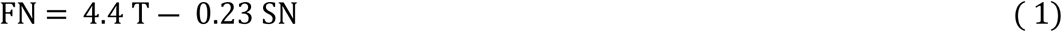

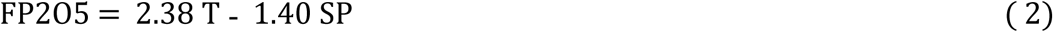

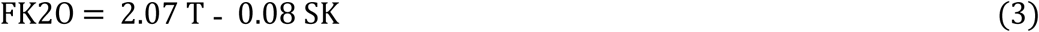

Where, FN = Nitrogen (kg ha^−1^) to be applied from fertilizer (1), FP_2_O_5_ = Phosphorus (kg ha^−1^) to be applied from fertilizer (2), FK_2_O = Potash (kg ha^−1^) to be applied from fertilizer (3), T = Targeted yield (t ha^−1^), SN = Available nitrogen (kg ha^−1^) from the soil, SP = Available phosphorus (kg ha^−1^) from the soil and SK = Available potassium (kg ha^−1^) from the soil. The fertilizer rates were reduced by 80% of the total amount of STCR based fertilizer dose because these equations were developed under optimum irrigation condition with 5 irrigations. For rainfed condition, maximum 3 irrigations can be provided and hence only 80% of the dose obtained from STCR equation is used. The biochar treatment was also kept as soil amendment (10 t ha^−1^) in combination of the T3 and T4 treatments. For the biochar preparation, drum method (indigenous technique) was used. Subabul (*Leucaena leucocephala*) biomass was collected from institute farm and sun dried. The subabul leaf, twigs and stem were chopped at desired size then pyrolized under oxygen-limited conditions and allowed to cool overnight. Subsequently, the biochar was crushed manually and ground to pass through a 5 mm sieve and applied in plot as per treatment details.. The chemical properties of original biochar are presented in Table 3. Biochar pH and electrical conductivity (EC) were determined in 1: 5 biochar to deionized water extraction [26]. Total C in original biomass and biochar sample were determined by dry combustion method using Shimadzu TOC analyzer. The ash alkalinity of biochar was determined by the method outlined by [27]. Briefly, the ash of biochar was obtained by heating the sample at 500°C for four hours in muffle furnace and then 0.5 g of biochar ash was dissolved in 25 ml of 1 *M* HCl. Subsequently, 5 ml of aliquot was titrated against 0.05*M* NaOH. Basic cations concentration (Ca^2+^, Mg^2+^, K^+^ and Na^+^) in aliquot was determined using the method as mentioned in case of soil. Biochar sample alkalinity was determined using the back titration method. In brief, 0.2 g of biochar sample was weighed into a beaker in triplicate followed by addition of 40 ml of 0.03*M* HCl solution. The samples were shaken for 2 hours on horizontal shaker at 25°C. After shaking samples were left undisturbed for 24 hours. Residual HCl was titrated to pH 7.0 with 0.05*M* NaOH. Biochar alkalinity was computed by determining the amount of HCl consumed.

**Table 3.** General characteristics of biochar used in the study

| Parameters | Value |
| --- | --- |
| pH(H<sub>2</sub>O) | 9.3 |
| EC (dS m<sup>-1</sup>) | 0.14 |
| Total C (g kg<sup>-1</sup>) | 79.9 |
| Ash (%) | 7.4% |
| Biochar alkalinity (cmol (p+) kg<sup>-1</sup>) | 39 |
| Ash alkalinity (cmol (p+) kg<sup>-1</sup>) | 580 |
| Exchangeable Ca<sup>2+</sup> (cmol (p+) kg<sup>-1</sup>) | 5.2 |
| Exchangeable Mg<sup>2+</sup> (cmol (p+) kg<sup>-1</sup>) | 3.3 |
| Exchangeable K<sup>+</sup> (cmol (p+) kg<sup>-1</sup>) | 17 |
| Exchangeable Na<sup>+</sup> (cmol (p+) kg<sup>-1</sup>) | 0.85 |

### Crop culture and management

The maize was planted during *kharif* or wet/rainy season on 10 July in 2014, 19 July in 2015 and 26 June in 2016 as rainfed crop. Before planting of the maize, field was leveled by tractor drawn laser leveler subsequently; two cross ploughings with disc plough and one ploughing with cultivator followed by planking were done for land leveling. Maize seed (Pro-agro 4212-95–110 day’s duration) was treated with imidacloprid @ 2g kg^−1^seed, to avoid the insect attack and the seed (20 kg ha^−1^) was planted at spacing at 0.60 m x 0.25 m. Glyphosate [N- (phosphonomethyl) glycine] was sprayed at 1.0 kg ha^−1^ in all plots about a week before maize planting for the controlling of weeds. Hand weedings were also done at 25 and 50 DAS (days after sowing) to check weed growth and at the time of second weeding, earthing up operation was also carried out. To avoid any early damage from insects, chloropyrifos (1.5 L ha^−1^) was sprayed a week after sowing.

### Growth and yield analysis

After the maize was planted to the field, 10 plants were tagged randomly in second row of north side. The data on different parameters under growth stage of maize i.e. knee high, tasseling, silking and physiological maturity stages in every treatment were recorded. For measurement of plant height (PH), dry matter accumulation (DMA) and leaf area index (LAI) the above-ground parts of the maize plants from 10 pre-determined plants from each treatment were sampled periodically over a knee high (V8), Tasseling (VT), silking (R1) and physiological maturity (PM). Plant height was measured from base of the stem to the base of last fully opened leaf periodically. Thereafter, plant samples taken to the laboratory and leaf area was measured from the leaves of sampled plants by using the leaf area meter (Model LI-COR-3100) and leaf area index was worked out as ratio of total leaf area to the land area of the sampled plants. Then, the plant samples were dried at 70 °C until constant weight and weighed to determine the dry matter weight. At maturity, pre-determined 20 plants of each treatment were used to measure the plant height, dry matter yield and yield attributes (excluding the border plants). The length of cobs was measured from the base to the tip of cob and girth of cobs was measured at three places *viz*. near the butt, in the middle and at the top with the help of a thread and measuring scale and the values thus obtained was averaged. The total number of kernel rows per cob and shelling percentage was also counted from selected cobs. While, total number of kernels in a row was counted from cobs selected for recording the number of grains grain row^−1^ and averaged out. The maize crop was harvested at maturity and sun dried at open place, then total biomass was weighed and recorded as total dry matter yield. The produce was then threshed and kernels were separated, dried (upto 12% moisture content) and weighed for grain yield. The maize grain yield was determined by adjusting the values with the corresponding shelling percentage and moisture conversion factor and the grain yield of all plots were finally presented uniformly at 12% moisture of three years data. Stover/ straw yield was obtained as difference between total dry matter yield and grain yield. The values were finally expressed in terms of t ha^−1^.

### Plant analysis for N accumulation and N use efficiency (NUEs)

Dry matter of whole plant, grain and stover samples of maize from each treatment were ground into powder by a grinding machine and 0.5 g dried powder were taken for digestion using Kjeldhal method. AR grade concentrated sulfuric acid was used for digestion of dried samples at temperature between 360 and 410 °C. The rate of digestion was accelerated by using copper sulphate as catalyst, and anhydrous sodium sulphate to raise the boiling temperature of sulfuric acid (H_2_SO_4_). After completion of digestion, the samples were cooled and diluted concentrated alkali (40% NaOH) was added to H_2_SO_4_ digest for distillation. The distilled ammonia was quantitatively absorbed in boric acid and titrated against standard acid (0.1 N H_2_ SO_4_) and the N content was calculated. Accordingly, N uptake pattern at different growth stages by maize was calculated by multiplying nutrient concentration with respective dry matter, accumulations/absorption. The agronomic efficiency (AE), partial factor productivity (PFP), physiological efficiency (PE) and recovery efficiency (RE) were calculated by the following formulas:

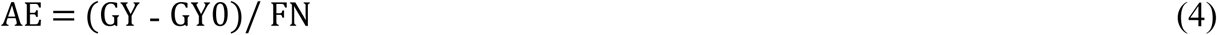

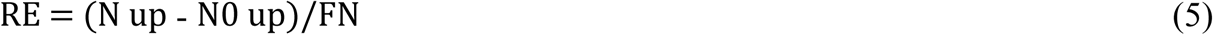

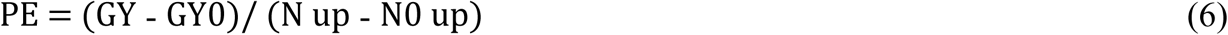

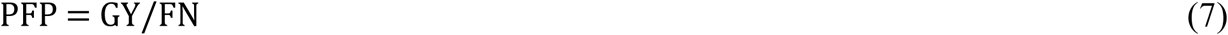

where GY0 and GY represent the grain yield in the N0 plot and fertilized N plots, respectively; and FN is the quantity of N fertilizer applied in N fertilized plot; N0 up and N up are the total nitrogen uptake in above ground biomass in the N0 plot and fertilized N plots, respectively.

### Soil Sampling and Analysis

Composite soil samples were collected from each plot using a core sampler from 0 to 15 cm depth after crop harvest. The samples were placed in plastic bags and transported to the laboratory, where air dry soil sample were homogenized, ground to pass through a 2 mm sieve and analyzed soil organic carbon (SOC) by the Walkley and Black method [28]. The concentration of ammonium (NH_4_^+^-N) and nitrate (NO_3_^-^-N) in soil were determined by shaking 5g soil with 50 ml of 2*M* KCl for 1 hour, followed by centrifugation and filtration through a filter paper. NH_4_^+^ and NO_3_^-^-N in filtered extracts were determined colorimetrically using flow injection analyzer (FOSS, UK).

### Statistical analysis

Data generated from the field experiments were subjected to the statistical analysis by using SAS 9.3 (SAS Institute, 2012). For combined analysis, treatment and year, both were treated as fixed effects in the linear model. Data from the years 2014, 2015 and 2016 were combined as the variances were homogeneous across these 3 years for all the parameters under study. Analysis of variance was performed using PROC GLM. The post hoc test for treatment mean comparison under each parameter was done on the basis of Tukey’s Honest Significant Difference (HSD) at *p*= 0.05 by using MEANS statement under PROC GLM with respect to the model error except year x treatment interaction when it was significant. Linear bivariate regression analysis was done between maize grain yield (as a dependent variable) and total N uptake (as an independent variable) using PROC REG. Factor analysis was conducted using PROC FACTOR to understand the factors that influence nitrogen uptake as well as to assess patterns or correlations among biomass and N accumulation variables. Factor analysis was conducted for two conditions namely N0 (N omission treatments) and N (treatments with N application). Variable factor maps were created using the first two factors obtained from the principal component analysis (PCA) method after varimax rotation. The correlation between two variables is represented by the cosine of the angle between them on variable factor map. Therefore, an acute angle implies positive correlation, an obtuse angle implies negative correlation, and a right angle denotes no correlation [29]. Principal component regression (PCR) between total dry matter yield with plant height, DMA, LAI, cob length, cob girth, 100 kernel weight, grains cob^−1^, N content and N uptake in dry matter was conducted using PROC PLS after resolving the multicollinearity problem among the independent variables. For detection of multicollinearity in the independent variables, correlation study was carried out using PROC CORR and for obtaining the optimum number of principal components; PCA was conducted with the independent variables using PROC PRINCOMP.

## Results

### Weather conditions

The meteorological data showed a marked variation in weather conditions during the three years of the experiment. Maximum and minimum temperatures remained almost constant during all the years, whereas, rainfall varied significantly. Maximum temperature ranged from 26.8-46.4 °C in 2014, 26.6-44.3 °C in 2015 and 25.3-45.0°C in 2016 with mean around 32.8 °C during the three years (Figs 1–3). Minimum temperature also varied from 15.1-31.9 °C in 2014, 14.5-30.9 °C in 2015 and 13.3-33.3°C in 2016 with mean around 23.3 °C during the three years. The pattern of rainfall was varied during the three years, in 2014, 89.3% (1009.5 mm) and in 2015, 87.3% (987.5 mm) of mean annual rainfall of 1130 (last 30 years) received during south-west monsoon period i.e. June to October. However, in 2016, crop received 1488 mm of rainfall which was 31.7% higher than the mean annual rainfall. A slight variation in grain yield and total biomass was due to fluctuation in weather condition and cumulative effect of growing degree days (GDD) in all three years. The GDD was lowest in 2014 while 2015 had the highest GDD during grain formation period.

### Plant height and leaf area index (LAI)

The maize plant height and LAI (Model LI-COR-3100) determined under different treatments in all three experimental years at different growth stages i.e. knee high (V8), tasseling (VT), silking (R1) and physiological maturity (PM) of maize (Table 4 and Fig 4). Pooled results indicated that the agronomic N management based on multi top dressing had significant effect on plant height and LAI. The identical trends among the different multi top dressing and biochar application was that the STCR based N dose (172.7 kg Nha^−1^) showed highest plant height and LAI values at all the V8, VT, R1 and PM stages and the absolute control (N0P0K0) and N control (N0) always had the smallest plant height and lowest LAI values. Among the different split-top dressing N doses, the plot which received full dose N (120 kg Nha^− 1^) into 3 unequal splits in combination of biochar (T11) produced tallest plants at V8 and VT and higher values of LAI at V8 only. However, at R1 and PM stages, the tallest plant and higher values of LAI were observed with treatments where the total amount of N rates (120 kg Nha^−1^) was applied into 2 equal splits at V8 and VT growth stages; T5, T7, T10, and T11 demonstrated comparable and non significant differences in the plant height of maize at all growth stages (Table 4).

**Table 4.** Comparison of the plant height under different agronomic N management practices and biochar application during crop growth stages (Pooled mean of 3 years)

| Treatment† | Plant height (cm) |  |  |  |
| --- | --- | --- | --- | --- |
|  | Days after planting |  |  | Physiological Maturity (PM) |
|  | Knee high (V8) | Tasseling (VT) | Silking stage (R1) |  |
| T1 | 28.2<sup>e</sup> | 64.3<sup>f</sup> | 81.6<sup>g</sup> | 122.8<sup>e</sup> |
| T2 | 29.7<sup>ef</sup> | 67.1<sup>f</sup> | 87.3<sup>g</sup> | 130.0<sup>e</sup> |
| T3 | 40.3<sup>bc</sup> | 87.8<sup>cd</sup> | 133.2<sup>cd</sup> | 163.9<sup>bcd</sup> |
| T4 | 38.3<sup>bc</sup> | 82.6<sup>de</sup> | 116.6<sup>ef</sup> | 158.0<sup>d</sup> |
| T5 | 32.8<sup>de</sup> | 95.1<sup>bc</sup> | 136.5<sup>bcd</sup> | 177.2<sup>a</sup> |
| T6 | 32.4<sup>def</sup> | 83.0<sup>de</sup> | 119.5<sup>e</sup> | 161.4<sup>cd</sup> |
| T7 | 30.1<sup>ef</sup> | 90.3<sup>cd</sup> | 138.0<sup>bc</sup> | 173.0<sup>abc</sup> |
| T8 | 28.9<sup>ef</sup> | 77.4<sup>e</sup> | 106.4<sup>f</sup> | 155.2<sup>d</sup> |
| T9 | 30.9<sup>ef</sup> | 95.2<sup>bc</sup> | 127.1<sup>d</sup> | 160.8<sup>d</sup> |
| T10 | 44.8<sup>a</sup> | 110.1<sup>a</sup> | 162.7<sup>a</sup> | 183.9<sup>a</sup> |
| T11 | 41.2<sup>ab</sup> | 100.0<sup>b</sup> | 146.4<sup>b</sup> | 174.4<sup>ab</sup> |
| T12 | 36.6<sup>cd</sup> | 87.9<sup>cd</sup> | 142.3<sup>bc</sup> | 164.1<sup>bcd</sup> |

Pooled means followed by the same letters are not significantly different according to Tukey's Honest Significant Difference (P=0.05)

**Fig. 4.**
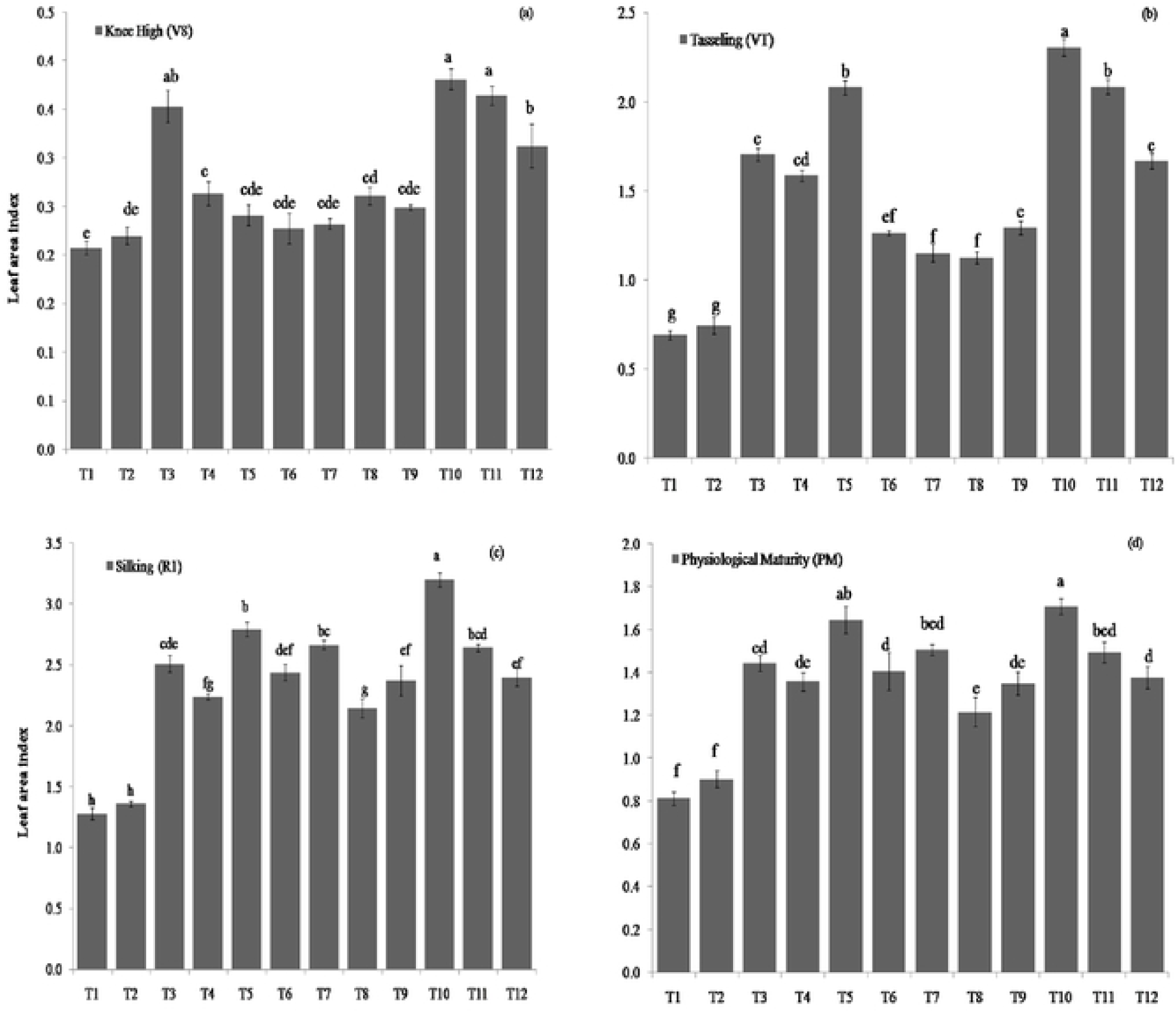
Effect of the treatment on leaf area index at knee high (V8), tasseling (VT), silking (R1) and physiological maturity (PM) of maize (pooled mean of three years data). Means bars marked by the same letters are not significantly different according to Tukey’s Honest Significant Difference (HSD) (P=0.05)

Furthermore, combined results showed that the application of biochar with 3 unequal split doses of N (120 kg Nha^−1^) improved the plant height at all growth stages. The skipping of basal N dose showed crop stress at initial growth stage (V8 and VT), but after receipt of N as top dressing at knee high (V8) and tasseling (VT), increased plant height and LAI values was observed. There were no significant difference in plant height and LAI at silking (R1) and physiological maturity (PM) when basal N rate skipped and total amount was applied in 3 equal splits. On other hand, the plant height and LAI increased significantly when total amount of N rate (120 kg Nha^−1^) was applied at V8 and VT as compared to lower N rate (90kg Nha^−1^) applied in 3 splits (120 kg Nha^−1^) at V8, VT and R1 stages.

### Biomass and nitrogen (N) accumulation

Combined analysis of the three years data indicated that the dry matter (g plant^−1^) significantly varied with different treatments, different growth stages viz., at the knee high (V8), tasseling (VT), silking (R1) and physiological maturity (PM) (Fig 5). The differences among the treatments were significant at different growth stages in all the three years, and highest biomass was observed with STCR based N application when compared to the remaining treatments. Among the different time and rates, the highest biomass accumulation was observed when, basal N rate was skipped and total amount of N was applied into two equal split (T5) at V8 and VT stages. However, the highest biomass production was recorded at PM stage. Application of biochar (10 t ha^−1^) with full amount of N fertilizer (T3) into 3 unequal splits were also significantly improved the biomass accumulation at V8, VT, R1 and PM stages of maize. As a similar fashion of N accumulation in dry matter, the N accumulation was the highest at PM followed by the R1, VT, and the lowest was in the V8 of maize growth stages (Figs 6 and 7). The highest N accumulation in biomass was recorded with the STCR based fertilization (T10). However, among the varying rates and time of application, the highest N accumulation in biomass was observed with the application of total amount of N (120 kg N ha^−1^) applied into 2 equal splits at V8 and VT growth stages while basal N rate was omitted (T5) as compared to other treatments. Remaining treatments were comparable and showed non-significant differences at all growth stage in respect to N accumulation in whole plant biomass during the experimentation.

**Fig. 5.**
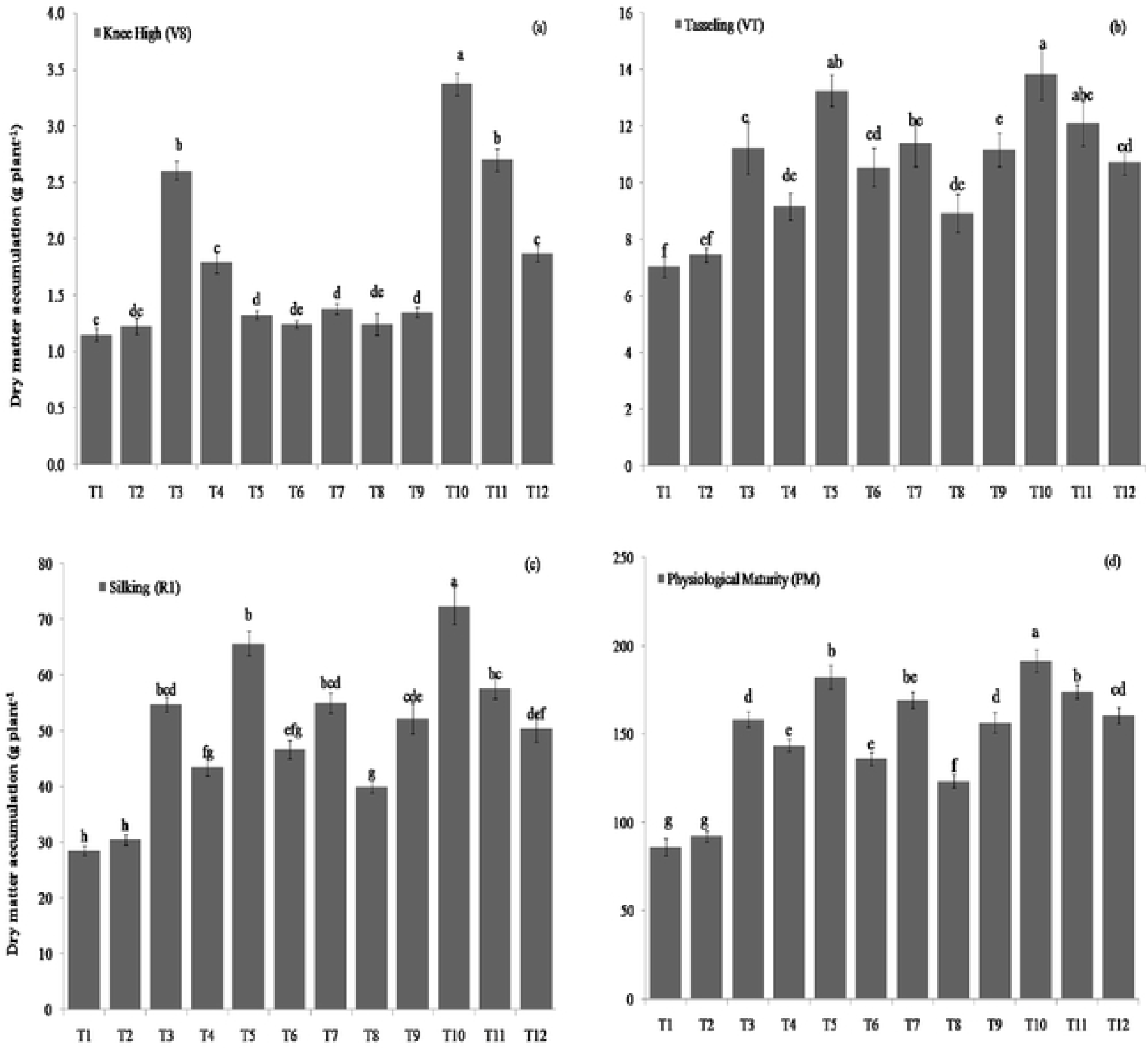
Effect of the treatment on dry matter accumulation at knee high (V8), tasseling (VT), silking (R1) and physiological maturity (PM) of maize (pooled mean of three years data). Means bars marked by the same letters are not significantly different according to Tukey’s Honest Significant Difference (HSD) (P=0.05).

**Fig. 6.**
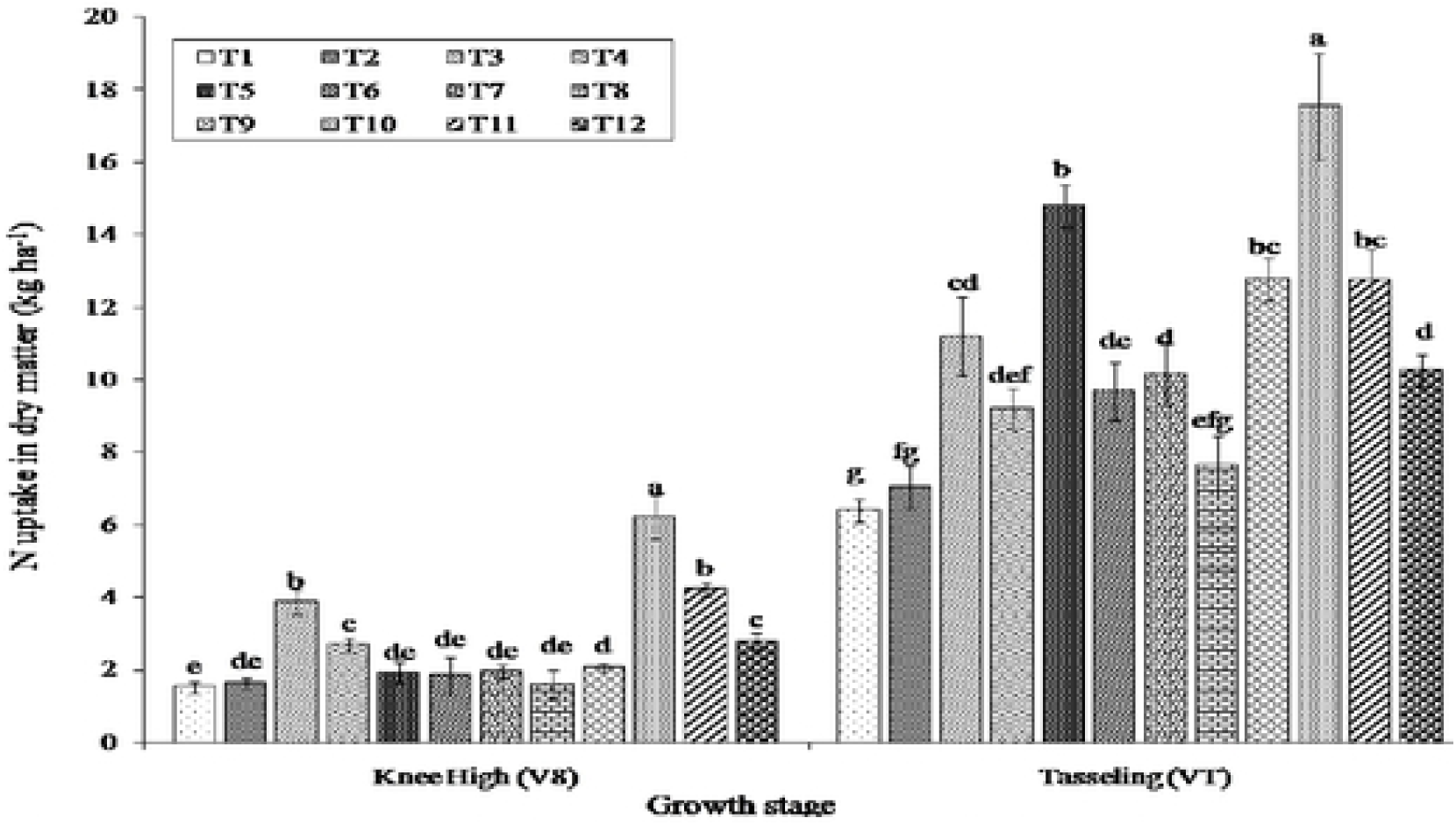
Effect of the treatment on N uptake in dry matter at knee high (V8) and tasseling (VT) stages of maize (pooled mean of three years data). Means bars marked by the same letters are not significantly different according to Tukey’s Honest Significant Difference (HSD) (P= 0.05).

**Fig. 7.**
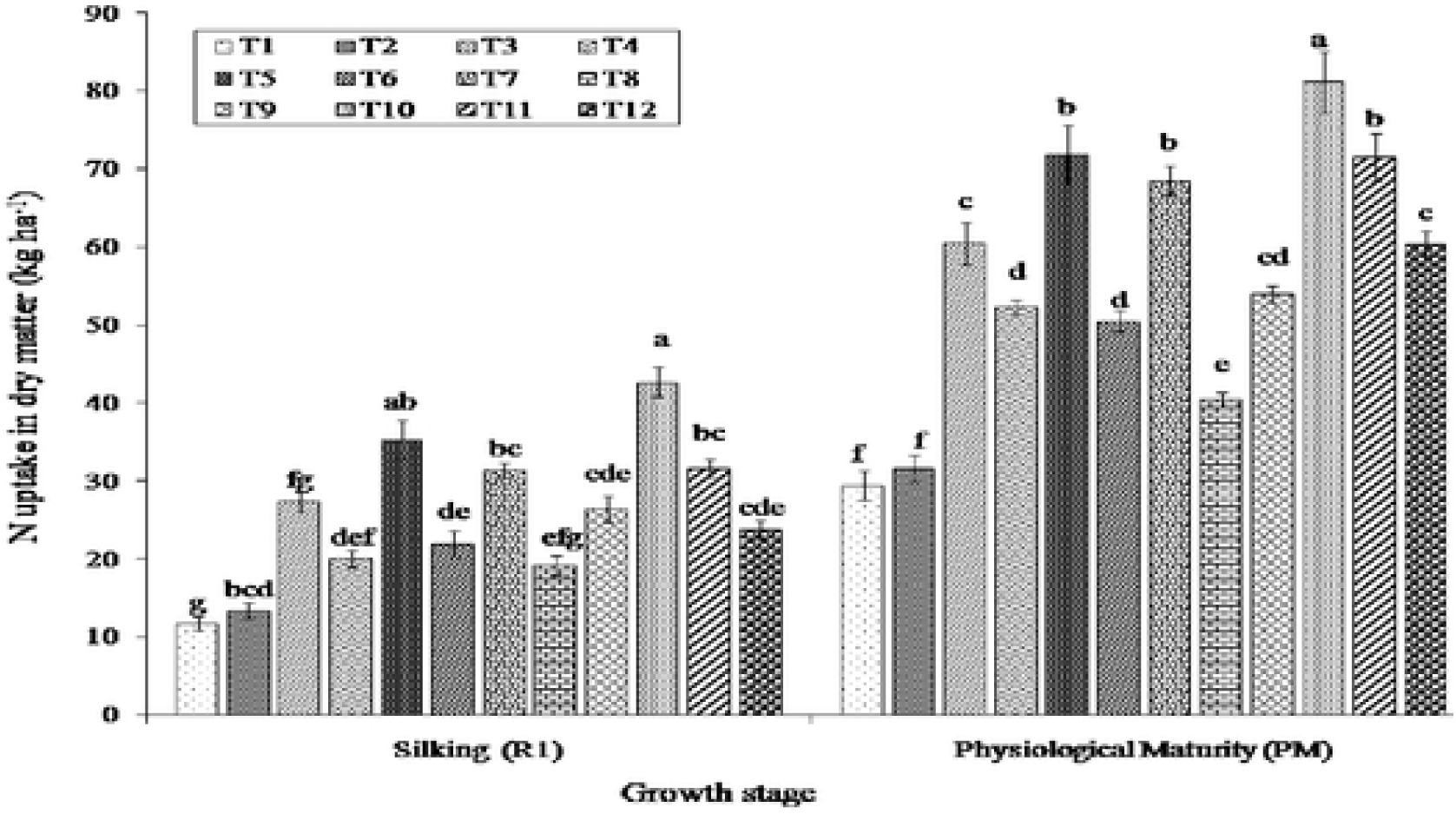
Effect of the treatment on N uptake in dry matter accumulation at silking (R1) and physiological maturity (PM) stgacs of maize (pooled mean of three years data). Means bars marked by the same letters are not significantly different according to Tukey’s Honest Significant Difference (HSD) (P=0.05).

### Maize yields and yield components

Pooled results indicated that the spliting of N rates had significant effect on yield components like cob length, cob girth, 100-kernel weight and number of grains cobs^−1^ during all three years (Table 5). The crop grown under STCR based NPK application exhibited the greatest yield components among all the treatments. However, among the different split top dressing, the higher values of yield attributing parameters were recorded with the application of N (120 kg Nha^−1^) in 2 equal splits at V8 and VT and omission of basal N.

**Table 5.** Comparison of the yield components under different agronomic N management practices and biochar application (pooled mean of 3 years)

| Treatment† | Yield components |  |  |  |
| --- | --- | --- | --- | --- |
|  | Cob length (cm) | Cob girth (cm) | 100 -grain weight (g) | No of grains cobs<sup>-1</sup> |
| T1 | 11.9<sup>f</sup> | 10.8<sup>e</sup> | 17.2<sup>b</sup> | 216.2<sup>h</sup> |
| T2 | 12.8<sup>f</sup> | 11.6<sup>de</sup> | 18.1<sup>b</sup> | 231.9<sup>h</sup> |
| T3 | 16.5<sup>cd</sup> | 14.0<sup>bc</sup> | 22.5<sup>a</sup> | 357.0<sup>de</sup> |
| T4 | 14.3<sup>e</sup> | 12.8<sup>cd</sup> | 19.2<sup>b</sup> | 313.5<sup>fg</sup> |
| T5 | 17.8<sup>b</sup> | 14.5<sup>ab</sup> | 22.0<sup>a</sup> | 425.6<sup>ab</sup> |
| T6 | 15.2<sup>e</sup> | 12.9<sup>cd</sup> | 19.1<sup>b</sup> | 331.3<sup>ef</sup> |
| T7 | 17.2<sup>bc</sup> | 14.7<sup>ab</sup> | 22.2<sup>a</sup> | 404.0<sup>bc</sup> |
| T8 | 14.4<sup>e</sup> | 12.4<sup>d</sup> | 18.8<sup>b</sup> | 284.3<sup>g</sup> |
| T9 | 15.3<sup>de</sup> | 12.8<sup>cd</sup> | 18.0<sup>b</sup> | 325.7<sup>ef</sup> |
| T10 | 19.2<sup>a</sup> | 15.7<sup>a</sup> | 23.3<sup>a</sup> | 458.7<sup>a</sup> |
| T11 | 16.8<sup>bc</sup> | 14.6<sup>ab</sup> | 22.5<sup>a</sup> | 389.9<sup>cd</sup> |
| T12 | 15.5<sup>de</sup> | 13.8<sup>bc</sup> | 19.3<sup>b</sup> | 337.6<sup>ef</sup> |

Pooled means followed by the same letters are not significantly different according to Tukey's Honest Significant Difference (P=0.05)

Biochar application also improved the yield components as compared to no application of biochar. Lower rate of N (90 Nha^−1^) had no significant effect on yield components when it was applied into either 2 or 3 equal splits. Among the different yield components, the differences in the 100-grain weight were not significant among the applied treatments. The highest number of grains cob^−1^ for the T5 and T11 was mainly due to the cob length and cob girth from 17.8 to 19.2 cm which was greater than the other treatments. Grain yield of maize varied in all three years with the applied treatments and yield reduction was noticed with N control (N0) and absolute control (N0P0K0) treatments relative to all other treatments (Table 6).

**Table 6.**
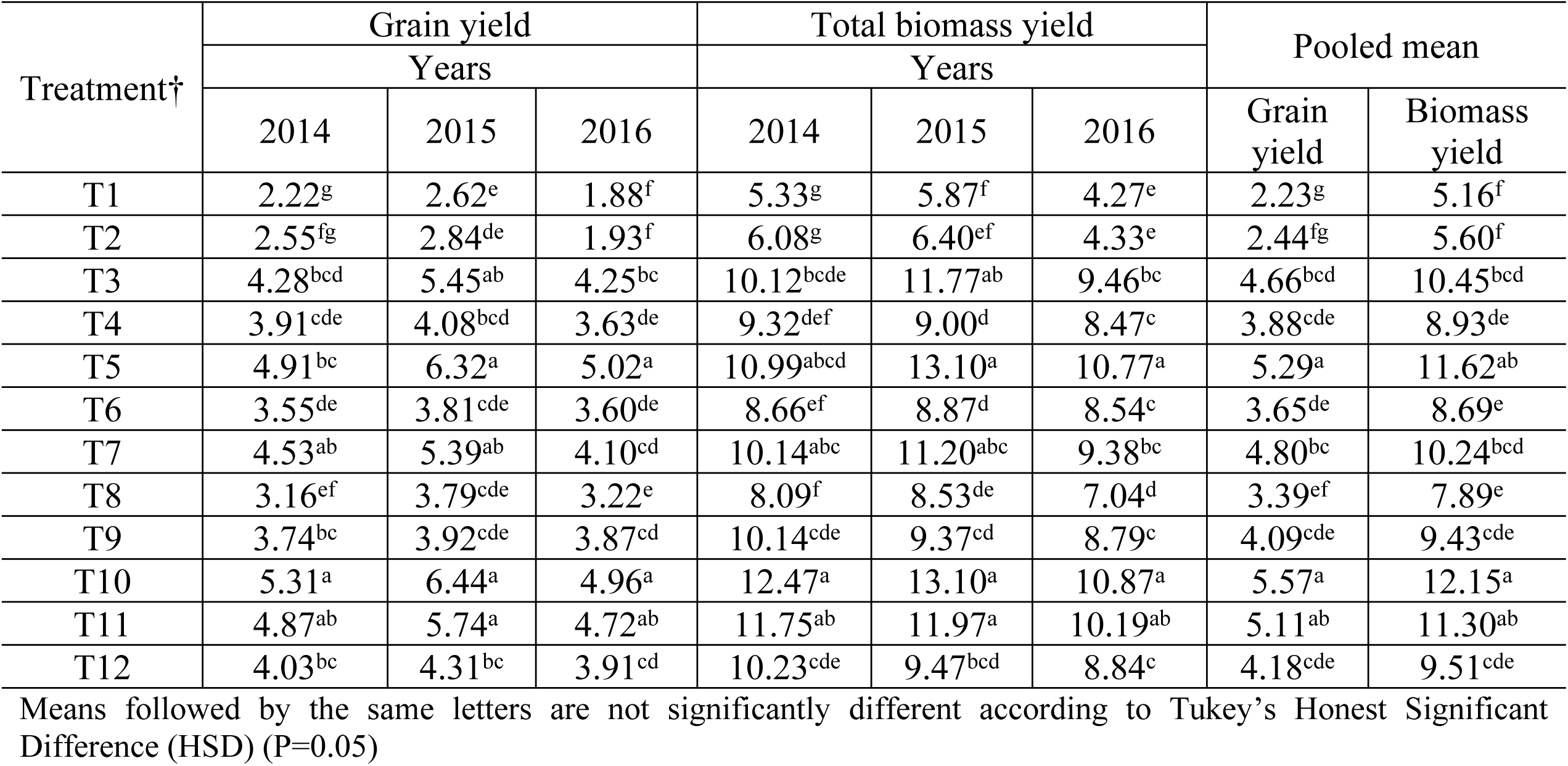
Comparison of the yield and biomass (t ha^−1^) of maize under the different agronomic N management practices and biochar application

STCR based NPK (T10) produced the maximum grain yield of 5.31, 6.44 and 4.96 tha^−1^ during 2014, 2015 and 2016, respectively. While, application of N (120 kg Nha^−1^) in 2 equal splits at V8 and VT with skipping of basal N rate (T5) recorded second highest grain yield of 4.91, 6.32 and 5.02 tha^−1^ during 2014, 2015 and 2016, respectively. However, the differences in grain yield between the T5 and the STCR were not found to be statistically significant except the first year (2014). In contrast, the grain yield of maize in the control and absolute control plots were consistently the lowest among all three years (Table 6). Furthermore, the grain yield was also significantly increased with the application of biochar (10 t ha^−1^) along with recommended dose of N into 3 splits as compared to no biochar application in all three years. STCR based NPK application recorded the highest total dry matter yield (TDMY) during all three years. Among the split application of N, the highest total biomass yield was obtained with the application of N (120 kg Nha^−1^) in 2 equal splits at V8 and VT with skipping of basal N rate (T5). Further, late split N application (T5) showed significant increases of 14.7%, 16.0% and 18.1% (with an average value of 16.3%) in grain yield than recommended doses (T3) in 2014, 2015 and 2016, respectively. With the combined analysis of 3 years’ data, highest grain yield (5.57 t ha^−1^) and TDMY (12.15 t ha^−1^) of maize were recorded with STCR NPK based fertilization (T10). While, among the varying N rate and time of application, grain yield (5.29 t ha^−1^) and TDMY (11.62 t ha^−1^) were significantly highest in the treatment where basal rate of N was skipped and total N (120 kg Nha^−1^) was applied in 2 equal splits (60 kg Nha^−1^) at V8 and VT, respectively (T5). This treatment (T5) increased grain yield and TDMY by 13.5% and 11.2% over recommended doses/rates, respectively. However, application of biochar along the recommended rate of N (120 kg Nha^−1^) was also significantly increased the grain yield and total biomass yield as compared to other treatments. Furthermore, the differences in grain yield and total biomass yield between the T5, T10 and T11 were not statistically significant.

### Nitrogen uptake and N use efficiencies

The total amount of N uptake in maize (grain+ straw) differed with the treatments. Combined result of N uptake was the highest with the STCR, and the lowest was in the N control (N0). However, among the split applications of N rates, the higher amount of N uptake was observed with the application of N rate (120 kg Nha^−1^) in 2 equal splits at V8 and VT with omission of basal N (T5) followed by the application of biochar (10 t ha^−1^) with recommended split as broadcast (T11). Evidently, grain yield of maize was highly dependent (*R*^*2*^=0.99 at p<0.0001) on N availability, as indicated by the fitted regression line of maize grain yield on N uptake (Fig 8.). The perfect positive linear relationship further illustrated that the change in grain yield is significantly associated with the changes in N uptake. Hence, the fact that, maximum N uptake was possible with the timely N application to the crop for higher productivity. Band placement of N (90 kg N ha^−1^) in three equal splits had no significant effect on total N uptake in all three years. Pooled data showed that the agronomic use efficiency of fertilizer N (AE) varied from the lowest of 11.0 kg kg^−1^ N to the highest of 28.3 0 kg kg^−1^ N among the different treatments (Table 7). By and large, the T5 demonstrated the greatest AE (28.3 kg kg^−1^ N), which was significantly greater than other treatments, except one treatment where the biochar was applied along with recommended split (T11). The plant treated with low rate of N (90 kg Nha^−1^) in 3 splits with skipping of basal dose always exhibited the lowest AE (11.0 kg kg^−1^ N), however, AE considerably increased with the application of biochar along with 90 kg Nha^−1^ (T12).

**Table 7.** Comparison of the N uptake, agronomic efficiency (AE), partial factor productivity (PFP), physiological efficiency (PE) and recovery efficiency (RE) under the different agronomic N management practices and biochar application (pooled of 3 years)

| Treatment<br>† | N uptake(kg ha<sup>-1</sup>) | AE (kg kg<sup>-1</sup> N) | PFP (kg kg<sup>-1</sup> N) | PE (kg kg<sup>-1</sup> N) | RE (%) |
| --- | --- | --- | --- | --- | --- |
| T1 | 48.3<sup>f</sup> | - | - | - | - |
| T2 | 52.3<sup>f</sup> | - | - | - | - |
| T3 | 98.6<sup>c</sup> | 19.5<sup>bc</sup> | 38.8<sup>cd</sup> | 47.7<sup>abc</sup> | 38.6<sup>bc</sup> |
| T4 | 83.6<sup>d</sup> | 17.3<sup>cd</sup> | 43.1<sup>abc</sup> | 46.0<sup>cd</sup> | 34.8<sup>bc</sup> |
| T5 | 109.9<sup>ab</sup> | 28.3<sup>a</sup> | 49.7<sup>a</sup> | 49.1<sup>ab</sup> | 48.0<sup>a</sup> |
| T6 | 80.4<sup>de</sup> | 13.6<sup>de</sup> | 37.3<sup>cde</sup> | 43.9<sup>de</sup> | 31.1<sup>cd</sup> |
| T7 | 103.2<sup>bc</sup> | 19.7<sup>bc</sup> | 39.0<sup>cd</sup> | 46.5<sup>bcd</sup> | 42.4<sup>ab</sup> |
| T8 | 74.0<sup>e</sup> | 11.0<sup>e</sup> | 34.8<sup>de</sup> | 42.0<sup>e</sup> | 24.1<sup>d</sup> |
| T9 | 87.7<sup>d</sup> | 16.9<sup>cd</sup> | 42.5<sup>bc</sup> | 45.9<sup>cd</sup> | 39.3<sup>bc</sup> |
| T10 | 114.6<sup>a</sup> | 18.7<sup>bcd</sup> | 31.9<sup>e</sup> | 50.2<sup>a</sup> | 35.6<sup>bc</sup> |
| T11 | 107.5<sup>abc</sup> | 23.3<sup>ab</sup> | 42.6<sup>bc</sup> | 48.6<sup>abc</sup> | 45.9<sup>a</sup> |
| T12 | 89.0<sup>d</sup> | 20.7<sup>bc</sup> | 46.5<sup>ab</sup> | 48.8<sup>abc</sup> | 40.7<sup>bc</sup> |

Pooled means followed by the same letters are not significantly different according to Tukey's Honest Significant Difference (P=0.05)

**Fig. 8.**
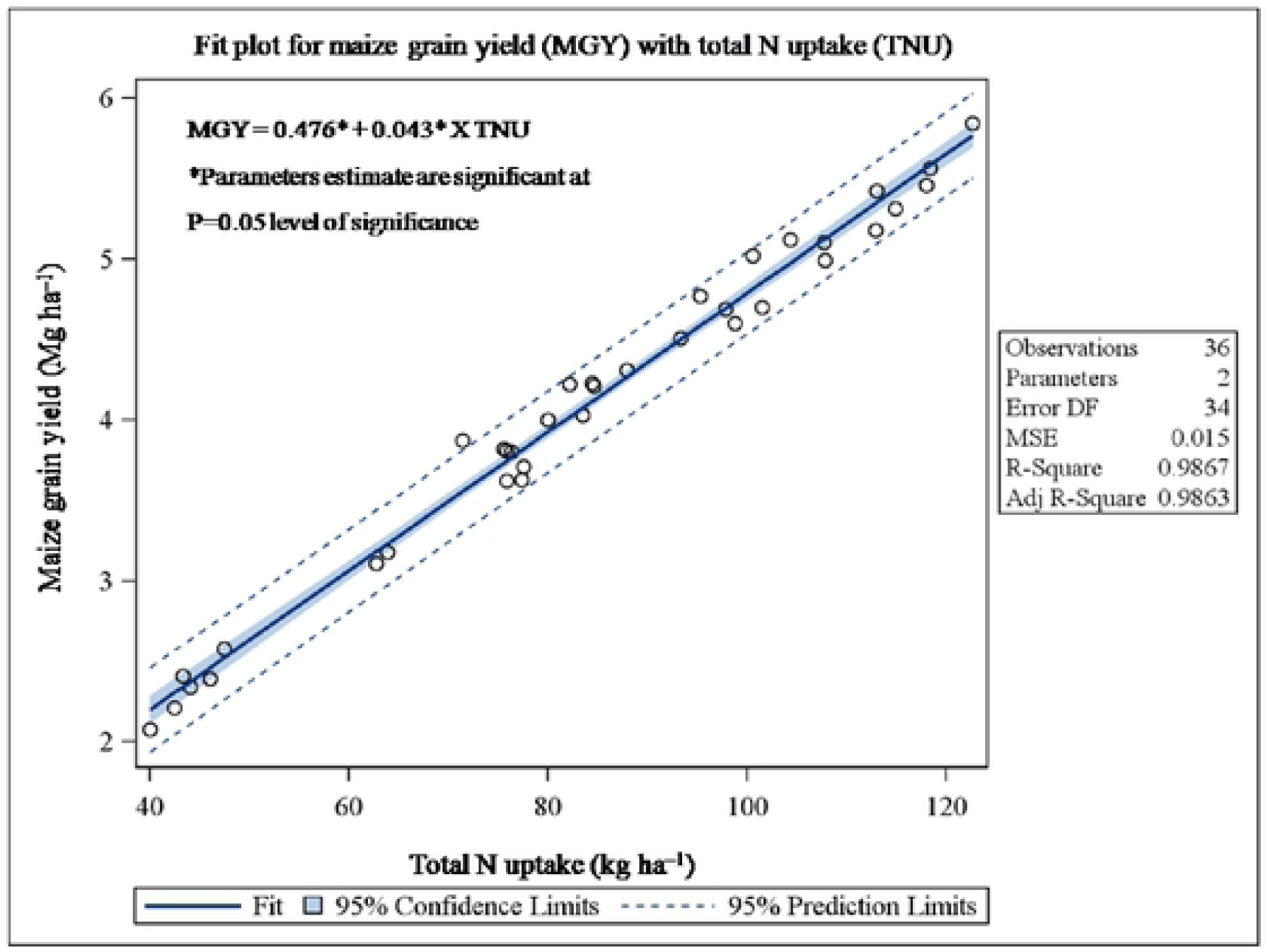
Relationship between total N uptake to maize grain yield under different agronomic N management practices

As a similar trend of AE, the application of N rate (120 kg Nha^−1^) in 2 equal splits at V8 and VT noticed the significant maximum PFP (49.7 kg kg^−1^ N) as compared to all treatments except T4 and T12. Furthermore, among the all treatments, STCR based N rate always exhibited the lowest PFP (31.9 kg kg^−1^ N). Band placement of N (90 kg Nha^−1^) along with biochar (10 t ha^−1^) also significantly increased the PFP in maize as compared to broadcast application (Table 7). Whereas, the difference in PE between different treatments were not significant, however, the application of N (120 kg N ha^−1^) into 2 or 3 split application significantly increased the PE as compared to N application equivalent to 90 kg N ha^−1^. The T5 recorded the highest RE (48.0%) in maize, which was significantly higher than other treatments, however application of biochar along with recommended split application of N (T11) was found non-significant. Furthermore, pooled analysis of 3 years’ data showed that the late multi-split N application at V8 and VT showed significant increases of 45.1%, 15.3%, 14.0% and 37.9% in AE, PFP, PE and RE than recommended practices (T3), respectively.

### Factors influencing N uptake and N use efficiency (NUEs)

Pooled analysis clearly indicated that the skipping of basal N rate and application of total amount of N applied at V8 and VT growth stages improved the N uptake and NUEs. Therefore, it is of interest to determine the factors of N uptake and NUEs to predict the maize grain yield that may increase the probability of response to multi-split N applications. Factor analysis was performed to understand the factors responsible for the relationship among the crop parameters with N content and N uptake in dry matter at different growth stages (V8, VT, R1 and PM) of rainfed maize. For factor analysis, the 16 variables were selected in the N treatments where as 13 variables were chosen for N0 treatments and AE, PE and RE were not included in N0 treatments due to no-application of N fertilizer.

In the N0 treatments (Fig 9A), N content and N uptake in biomass had high significant correlation with DMA at V8 (r=0.65 and 0.97), VT (r=0.74 and 0.97), R1 (NUDM; r=0.84) and PM (NUDM; r=0.84) growth stages with significant poor relation with DMA at PM (NDM; r=0.50). It also revealed that there was no significant relation between Nitrogen content in dry matter (NDM) and DMA at R1 as the angle between the vectors of NDM and DMA at R1 was approximately measured by a right angle (Kroonenberg, 1995) on variable factor map (Fig. 9A). On the other hand, there was also no significant influence of N content and N uptake in biomass on TDMY at PM growth stage under N0 treatments.

**Fig. 9.**
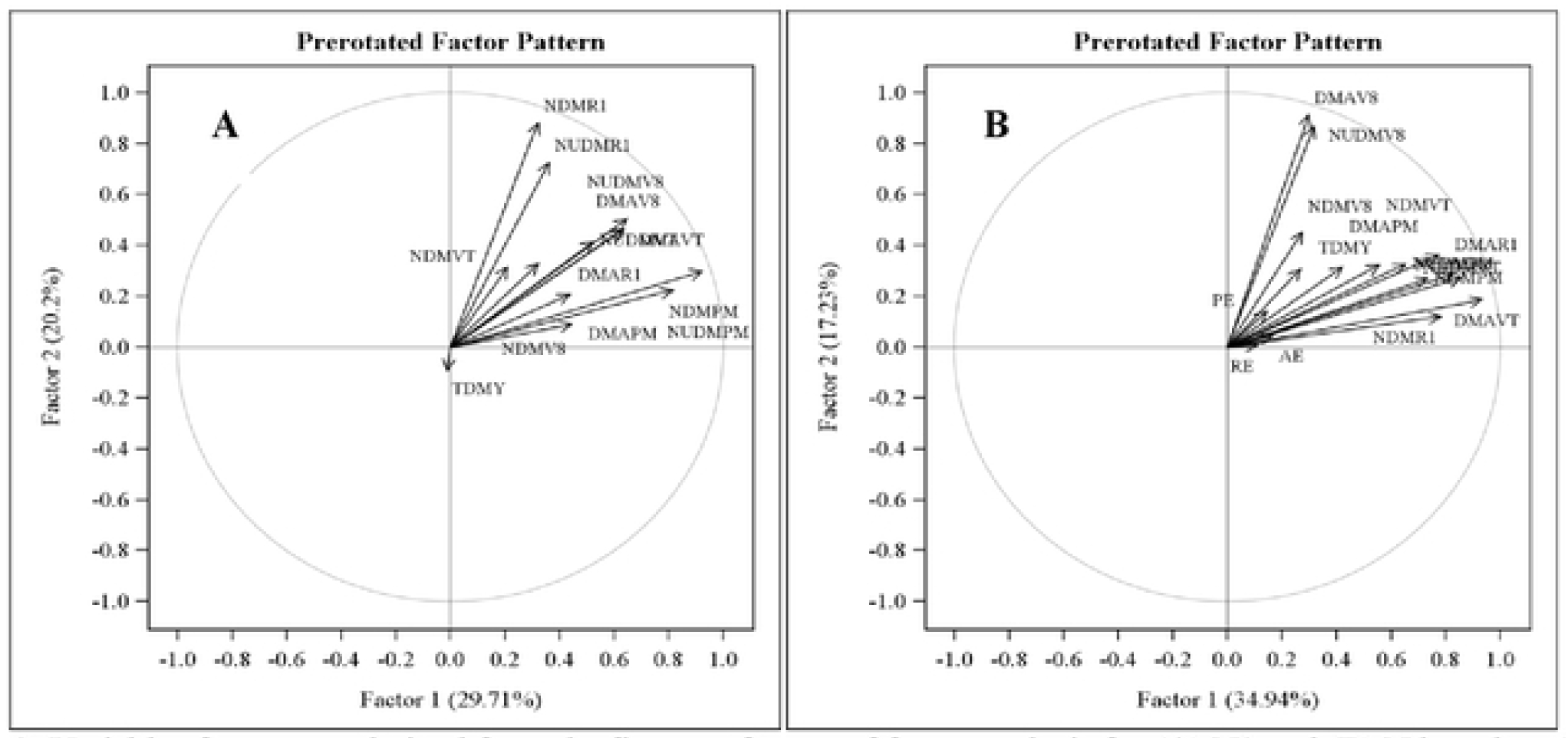
Variables factor map derived from the first two factors of factor analysis for (A) N0 and (B) N based on 3 years data (2014, 2015 and 2016). DMAV8/NDMV8/NUDMV8, dry matter accumulation (kg ha^−1^)/N content (%) in dry matter/ N uptake in dry matter (kg ha^−1^)at V8 growth stage; DMAVT/NDMVT/NUDMVT, dry matter accumulation /N content in dry matter/ N uptake in dry matter at VT growth stage; DMAR1/NDMR1/NUDMR1, dry matter accumulation/N content in dry matter/ N uptake in dry matter at R1 growth stage; DMAPM/NDM PM/NUDM PM, dry matter accumulation/N content in dry matter/ N uptake in dry matter at PM growth stage; TDMY, total dry matter yield (kg ha^−1^); AE, agronomic efficiency (kg kg^−1^ N); PE, physiology efficiency (kg kg^−1^ N); RE, recovery efficiency (%).

Under N treatments (Fig 9B), N content and N uptake in biomass had strong significant correlation with DMA at V8 (r=0.64 and 0.98), VT (r=0.71 and 0.96), R1 (r=0.76 and 0.96) and PM (r=0.62 and 0.89) growth stages. Furthermore, results indicated that DMA was most closely related to AE (r=0.55) whereas poorly related to PE (r=0.37) and RE (r=0.48) at PM growth stage. Total dry matter yield (TDMY) was also highly associated with, NUDM (r=0.72), AE (r=0.58) and RE (r=0.52), however, meagerly related to NDM (r=0.47) and PE (r=0.33). Moreover, the study showed the positive and significant linear association of dry matter accumulation (biomass) with N uptake in dry matter at V8, VT, R1 and PM growth stages of maize. The relationship between biomass and N uptake in biomass turned out to be more robust at V8 followed by VT, R1, and PM over the years.

### Principal component regression (PCR) relationship to total dry matter yield (TDMY)

Principal component analysis (PCA) was used to identify the factors influencing total dry matter of maize yield and has been used as the preliminary step in the development of a prediction model for TDMY as the Pearson correlation analysis revealed the result that the independent variables were significantly correlated to each other (Table 8) i.e. the multicollinearity was present among the independent variables.

**Table 8.** Correlation coefficient (n = 90) with exact probability level of significance for independent variables†

| Pearsons' Correlation Coefficient | PH | LAI | DMA | CobL | CobG | GW | GrainC | NDM | NUDM |
| --- | --- | --- | --- | --- | --- | --- | --- | --- | --- |
| PH | 1 | 0.474<br>($<.0001$) | 0.625<br>($<.0001$) | 0.549<br>($<.0001$) | 0.573<br>($<.0001$) | 0.583<br>($<.0001$) | 0.732<br>($<.0001$) | 0.291<br>(0.0054) | 0.518<br>($<.0001$) |
| LAI | | 1 | 0.678<br>($<.0001$) | 0.711<br>($<.0001$) | 0.627<br>($<.0001$) | 0.541<br>($<.0001$) | 0.757<br>($<.0001$) | 0.709<br>($<.0001$) | 0.762<br>($<.0001$) |
| DMA | | | 1 | 0.788<br>($<.0001$) | 0.732<br>($<.0001$) | 0.633<br>($<.0001$) | 0.809<br>($<.0001$) | 0.621<br>($<.0001$) | 0.892<br>($<.0001$) |
| CobL | | | | 1 | 0.705<br>($<.0001$) | 0.659<br>($<.0001$) | 0.809<br>($<.0001$) | 0.576<br>($<.0001$) | 0.760<br>($<.0001$) |
| CobG | | | | | 1 | 0.608<br>($<.0001$) | 0.707<br>($<.0001$) | 0.493<br>($<.0001$) | 0.678<br>($<.0001$) |
| GW | | | | | | 1 | 0.670<br>($<.0001$) | 0.496<br>($<.0001$) | 0.633<br>($<.0001$) |
| GrainC | | | | | | | 1 | 0.576<br>($<.0001$) | 0.766<br>($<.0001$) |
| NDM | | | | | | | | 1 | 0.902<br>($<.0001$) |
| NUDM |  |  |  |  |  |  |  |  | 1 |

The values in the parentheses are the actual probability of significance (probability $> |r|$ under $H_0: R_{ho}=0$ )
†PH, plant height; LAI, leaf area index; DMA, dry matter accumulation; CobL, cob length; CobG, cob girth; GW, 100 grain weight; GrainC, grains cob<sup>-1</sup>; NDM, N content in dry matter; NUDM, N uptake in dry matter.

To resolve the multicollinearity problem, principal component regression model was fitted for total dry matter yield (TDMY) with respect to the independent variables *viz*., plant height (PH); LAI, DMA, cob length (CobL), cob girth (CobG), 100-grain weight, grains cob^−1^ (grainC), N content in dry matter (NDM), N uptake in dry matter (NUDM). In the analysis, 3 principal components were selected because these components explained the cumulative variation of 99.22% in independent variables data set (Fig 10). PCA as illustrated from the PC loading coefficient of independent variables revealed that first PC (PC1) contributed 92.98 % explained variation (Fig 10) and dominated by grains cob^−1^, LAI and NUDM (Table 9).

**Table 9.** Principal Component (PC) loading coefficient for each independent variable

| PCs | Plant height | Leaf area index | Dry matter accumulation | Cob length | Cob girth | 100-grain weight | grains cob<sup>-1</sup> | N content in dry matter | N uptake in dry matter |
| --- | --- | --- | --- | --- | --- | --- | --- | --- | --- |
| PC1 | 0.1457 | 0.2904 | 0.0022 | 0.0233 | 0.0156 | 0.0261 | 0.9255 | 0.0007 | 0.1906 |
| PC2 | -0.0614 | 0.7687 | 0.0016 | 0.0225 | 0.0202 | 0.0204 | -0.3432 | 0.0023 | 0.5349 |
| PC3 | 0.9512 | 0.1549 | -0.0036 | -0.0164 | 0.0117 | 0.0303 | -0.1550 | -0.0028 | -0.2136 |

**Fig. 10.**
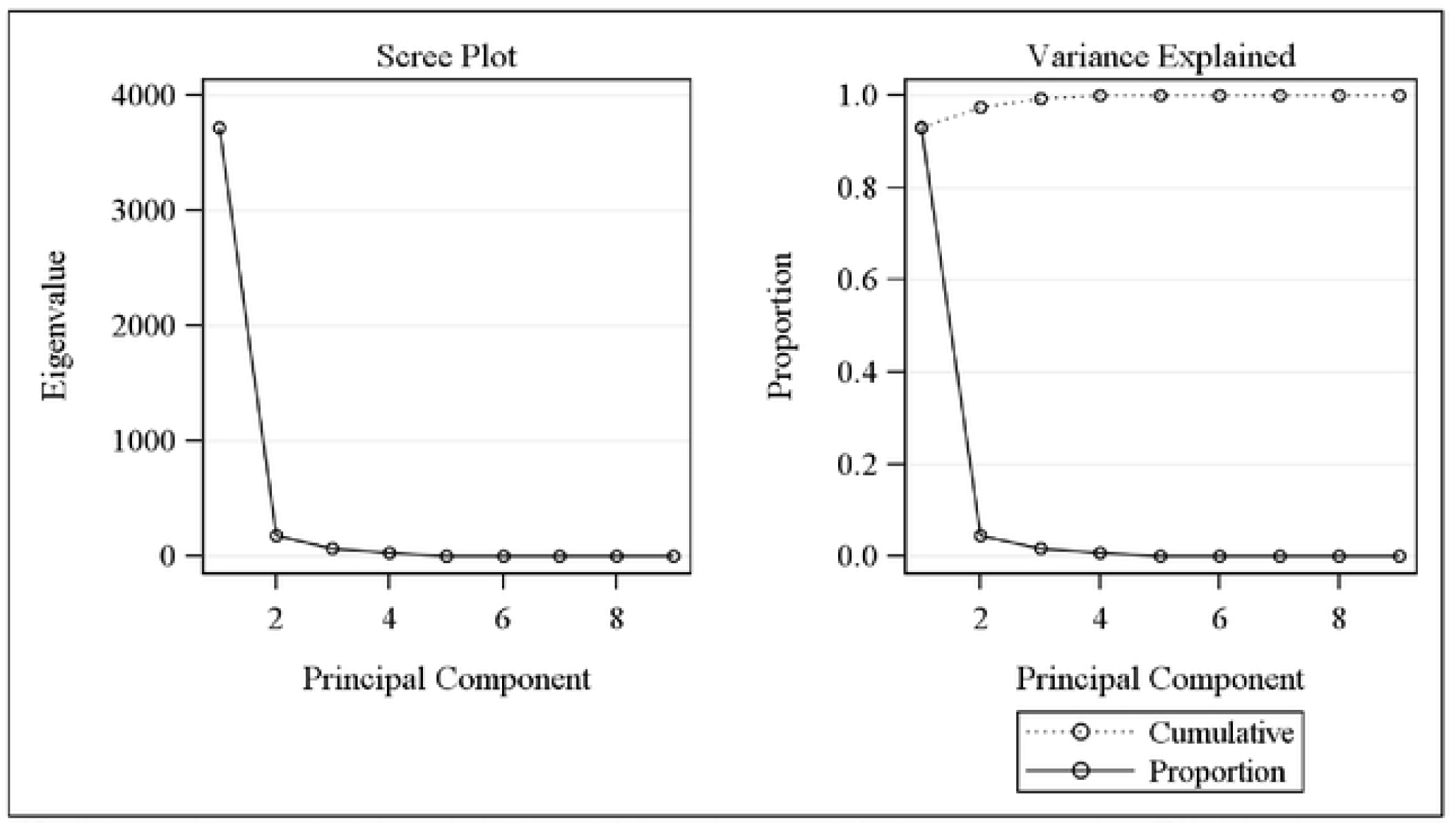
Scree and variance explained plot derived from the principal component anaylsis (PCA) for detection of the optimum number of principal components

The second and third PCs (PC2 and PC3) explained an additional 4.55% and 1.69% of the variation and was predominated by grains cob^−1^ and NUDM, respectively. The fitted principal component regression model based on these 3 PCs had explained the cumulative variation of 84.95% in model effects and of 78.45 % in dependent variable (TDMY) (Table 10).

**Table 10.** Percent variation accounted for by Principal Components (PCs) in Principal Component Regression (PCR) for total dry matter yield with independent variables†

| Number of extracted factors | Model effects |  | Dependent variable |  |
| --- | --- | --- | --- | --- |
|  | Current | Total | Current | Total |
| PC 1 | 69.99 | 69.99 | 69.69 | 69.69 |
| PC 2 | 10.11 | 80.10 | 7.52 | 77.21 |
| PC 3 | 4.85 | 84.95 | 1.24 | 78.45 |

† Plant height, leaf area index, dry matter accumulation, cob length, cob girth, 100 grain weight, grains cob<sup>-1</sup>, N content and N uptake in dry matter.

The fitted principal component regression model with significant parameter estimate corresponding to the independent variable for prediction of TDMY was given in equation (8).

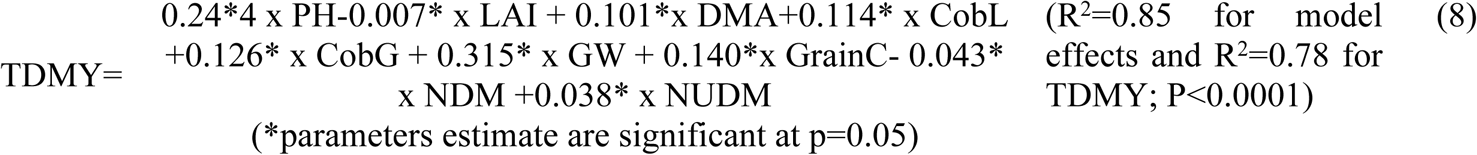

### Soil Chemical properties

There was an improvement in the soil chemical properties under the different agronomic management practices at the end of the study (Table 11). Application of N along with biochar significantly increased the soil organic carbon (SOC), concentration of ammonium (NH_4_^+^-N) and nitrate (NO_3_^-^-N) in soil as compared to recommended agronomic practices. A perusal of data showed that the highest SOC (5.47 g kg^−1^) was recorded with application of N along with biochar at 10 t ha^−1^ (T11), while lowest SOC was noticed under the control treatment.

**Table 11.** Effect of different agronomic N management practices and biochar application on chemical properties of soil (0-15cm soil layer) at the end of study (2016–2017)

| Treatment | Soil organic carbon (SOC) (g $\text{kg}^{-1}$) | $\text{NH}_4\text{-N}$ (mg $\text{kg}^{-1}$) | $\text{NO}_3\text{-N}$ (mg $\text{kg}^{-1}$) |
| --- | --- | --- | --- |
| T1 | 4.03<sup>c</sup> | 1.19<sup>h</sup> | 0.21<sup>d</sup> |
| T2 | 4.17<sup>b</sup> | 1.56<sup>g</sup> | 0.22<sup>d</sup> |
| T3 | 4.50<sup>ab</sup> | 1.94<sup>def</sup> | 0.34<sup>b</sup> |
| T4 | 4.20<sup>ab</sup> | 1.78<sup>fg</sup> | 0.26<sup>bcd</sup> |
| T5 | 4.43<sup>ab</sup> | 2.30<sup>abc</sup> | 0.44<sup>a</sup> |
| T6 | 4.13<sup>b</sup> | 1.89<sup>ef</sup> | 0.29<sup>bcd</sup> |
| T7 | 4.43<sup>ab</sup> | 2.08<sup>cde</sup> | 0.50<sup>a</sup> |
| T8 | 4.17<sup>b</sup> | 1.82<sup>ef</sup> | 0.32<sup>bc</sup> |
| T9 | 4.23<sup>ab</sup> | 1.80<sup>fg</sup> | 0.24<sup>cd</sup> |
| T10 | 4.63<sup>ab</sup> | 2.50<sup>a</sup> | 0.48<sup>a</sup> |
| T11 | 5.47<sup>a</sup> | 2.41<sup>ab</sup> | 0.52<sup>a</sup> |
| T12 | 5.17<sup>ab</sup> | 2.16<sup>bcd</sup> | 0.50<sup>a</sup> |

Means followed by the same letters are not significantly different according to Tukey's Honest Significant Difference (P=0.05)

Further results showed that, there were no significant differences among the different split N application in relation to SOC at the end of crop season however; application of biochar improved the SOC (5.47g kg^−1^) level as compared non-biochar treatments. Application of biochar along with chemical fertilizer (120 kg Nha^−1^) significantly increased the concentration of ammonium (2.40 mg kg^−1^) and nitrate (0.52 mg kg^−1^) in soil (P<0.05) as compared to non-biochar treatments. Since, application of biochar causes amelioration effects, which results into improvement soil fertility. Furthermore, results showed that the application of N rate (120 kg Nha^−1^) in 2 equal splits at V8 and VT growth stages also recorded higher values of NH_4_^+^-N (2.29 mg kg^−1^) and NO_3_^-^-N (0.44 mg kg^−1^) concentration as compared recommended dose of N and control treatment. The application of STCR based N dose also increased the concentration of NH_4_^+^-N and NO_3_^-^-N in soil as compared to traditional fertilizer application and control treatment but this was due to heavy N fertilization.

## Discussion

### Growth attributes

Combined results indicated that the STCR based N dose had highest plant height and LAI values at all growth stages, probably due to high supply of N (172.7 kg Nha^−1^) sufficed for adequate growth of the crop. The role of N in the stimulation of cell division may have led to more leaf numbers and expansion development during the grand growth periods [30]. However, the fertilizer N response to crops mainly depends on soil N, crop demand and climatic conditions [18,31]. However, among the multi-split N application, N supplied into 3 unequal splits in the biochar treated plots produced tallest plants and higher values of LAI at knee high (V8) and Tasseling (VT) stages. This indicated that contributing effect of N on plant height and LAI can be explained on the basis of the fact that split N application at the early (V8) and mid grand (VT) growth stages enhanced the meristematic cells and their growth leading to increase in the leaf expansion and the formation of tassels [32]). Moreover, N also acts as a key constituent of all the amino acids in plant structure, chlorophylls, enzymes, protein, purines and pyrimidines which are essential for the growth and development of plant tissues [33]. Such result indicated that increase in plant height and LAI with split application of N fertilizer might be due to the fact that N boost plant growth, increases the number of nodes and internodes which results in progressive increase in the plant height and LAI [34]. Further, it was noted that with the skipping of basal N rate considering residual N in soil, although plant showed some minor stress in the beginning, but subsequent to application of 50% N at V8 increased the whole plant growth rapidly ([35]. When N is limiting in soil, N application rates may increase total N accumulation in plant which might have more stimulated vigorous growth of whole plant [19,36]. In contrast, the very high rate of N application used initially could not turn out higher growth attributes but possibly led to considerable N loss due to remobilization of N uptake by plants [37].

### Biomass and N accumulation

Biomass in terms of dry matter production and N accumulation responded positively to N rate at different growth stages. STCR produced highest biomass production primarily due to addition of higher rate of N which might have enhanced greater N accumulation in whole plant part stimulated the development of sink capacity and thereby resulted into greater biomass production [38]. The second highest biomass accumulation when the total amount of N (120 kg N ha^−1^) was applied into two equal splits at V8 and VT growth stages might have arisen due to more partitioning and assimilation to the stem and tassels that resulted in higher biomass [39]. In general improving crop yield and NUEs for maize could be achieved by determining the plant N status during the growing period. The N accumulation by maize was lowest at early growth stage (V8) and increased from VT and recorded highest at PM growth stage. This indicated that only small amount of N are required during early growth stage and the high concentration of N in the root zone is sufficient for promoting early growth [24]. Besides, N application at V8 and VT growth stages demonstrated stronger biomass accumulation ability [40] due to most favorable synchronization between time of N application and crop demand [41]. The N accumulation was significantly highest and strongly correlated with the biomass at different growth stages. This indicated that a greater N accumulation in biomass stimulated the higher plant growth with higher dry matter partitioning in maize [42]. Low N uptake in early growth stage might be due to excessive N fertilization under the conventional fertilization [34]. Application of biochar (10 t ha^−1^) in combination of multi split-top dressed N also increased the dry matter partitioning and N uptake which clearly demonstrated the positive impact of biochar in maize during all three year of the experimentation. This might be due to transient flush of labile compound into the rhizosphere that can enhance nutrient cycling and increase crop biomass and N accumulation in whole plant [43].

### Maize yields and yield components

The 3 years research results confirmed that skipping of basal N rate and late season N application can be effective in recording higher grain yield and yield components like cob length, cob girth, 100-kernel weight and number of grains cobs^−1^ during all three years. Late season N application had significant and positive correlation with the higher grain yield [44]. The N application at V8 and VT vegetative stages resulted in a significant grain yield as compared to other treatments. Therefore, a positive yield response to late split N application might be more synchronized with crop peak N demand compared to the basal N application [45]. Moreover, our hypothesis was that the native soil N is sufficient for the completion of N demand by the plant at early growth stage and a larger impact of N availability on grain yield was noticed when very severe N deficiencies occur during early growth period [19]. Wang et al. [46] were also reported the similar results that there is no negative impact on maize due to delaying the total amount of N until V11 growth stage. In contrast, N supplied at the time of planting (basal N rate), plant shows inability to accumulate the applied N during the early growth stages which might have led to great loss of large amount of basal N application to the environment due to poor root system under rainy season [47]. Similarly, some previous results also indicated that the higher N application (2-3 splits) at early growth stages resulted in a increase of soil ammonia concentration peak, as a result, the soil N would increase the loss due to leading to leaching or denitrification [48]. Moreover, there was no significant effect on maize yield response to late season split N application at silking (R1), which might be due to the fact that the maize plant was already N adequate [49]. The STCR based NPK demonstrated the greatest yield due to addition of more N which might have increased the cob length and girth and greater number of kernels cob^−1^. There were no yield differences among the treatments with split N applications at lower N rate (90 Nha^−1^) and no yield advantages for late split application except in biochar application. Grain yield of rainfed maize varied in all three years primarily due to large variation in rainfall pattern.

### Nitrogen uptake and N use efficiencies (NUEs)

In the recent study, nitrogen uptake and NUEs was significantly influenced by the multi-split N application due to better synchronization of N availability to crop peak demand of N. Although N uptake and NUEs depends on agronomic management practices, type of soil and climate factors ([50], combined analysis showed that the skipping of basal N rate and delayed application of total amount of N at V8 and VT significantly improved the AE, PFP, PE and RE in maize as compared to all treatments during all 3 years. This could be attributed due to more N accumulation during the post V8 and VT which led to higher total N uptake and resulted in increased NUEs in maize [51]. STCR based N rate always exhibited the lowest AE, PFP and RE which might be due to negative impact of more fertilization on grain yield, environmental quality and reduced NUEs. Therefore, there is a need to provide sufficient N fertilizer for maize [52,53]. Band placement of N along with biochar also improved the PFP in maize as compared to broadcast application. This indicated that the application of N through band placement is attributed to higher PFP due to full utilization of applied N, which might have positive impact on grain yield. Besides, AE, PFP, PE and RE were enhanced due to incorporation of biochar. This is attributed to the beneficial effect of biochar in soil N dynamics through altering soil chemical and biological properties [54] and biochar also influenced the N cycle of agro-ecosystems [55].

### Factors influencing N uptake and NUEs

Factor analysis was carried out to assess the factors responsible for the relations among the biomass and TDMY with N uptake, AE, PE and RE. It was reported that several factors make significant contribution to the N uptake and NUEs in maize [46]. Similarly, N contributes an important role in crop production particularly when it was applied at different timings and rates. Our 3 years results have consistent shown that N was the main limiting management factors of the poor yield and low NUEs [56]. In contrast, total amount of N applied at V8 and VT stages, improved the N uptake and NUEs (AE, PE and RE) in rainfed maize, which has the potential to increase the maize yield through minimizing N losses [57]. N uptake was strongly correlated with plant biomass at V8 and VT as compared to PM growth stage. This indicated that the V8 and VT were the critical growth stages for the N application for higher biomass production, which might be due to higher N uptake [58]. Furthermore, there was no influence of DMA on N content at R1 stage as indicated by variable factor map [59]. This might be explained by the poor synchrony between crop supply-demand and N uptake. Thus, N uptake was greatest predictor of TDMY and AE was most strongly related to DMA as compared to PE and RE. It also indicated that higher N uptake resulted in higher growth and yield as indicated by strong correlation [60]. Similarly, Nyiraneza et al.[61] also noticed that N availability in biomass indices might be able to explain the variation in maize yield and N uptake. Moreover, the study showed the positive and significant linear association of DMA (biomass) with N uptake at V8 and VT growth stages of maize. In general the relationship between biomass and N uptake turned out to be more robust to late split N-applications at V8 and VT growth stages over the years in rainfed maize.

### Principal component regression (PCR) relationship to total dry matter yield (TDMY)

The total dry matter yield (TDMY) was greatly influenced by the growth and yield attributes along with N content and uptake as illustrated by the principal component regression (PCR) model. Results obtained from the fitted PCR model showed that the cumulative variability of 78.45% in TDMY was explained by the retained three PCs used for parameter estimation. Model demonstrated based on the regression coefficient parameter estimated that GW (grain weight) act as strong positive predictor for TDMY followed by PH, GrainC, CobG, CobL, DMA and NUDM where as LAI and NDM had significant adverse influence on yield [58]. The validity of PCR model depends on the presence of multicollinearity among the independent variables (PH, LAI, DMA, CobL, CobG, GW, GrainC, NDM and NUDM). In addition, using PCs for resolving multicollinearity and the three optimum PCs which contributed 99.22% explained variation in independent variables data set acts as strength for PCR model for the prediction of TDMY. Similar finding were also observed by Ciampitti and Vyn [62]. It was showed that PC1 was explained the 69.69% variability in TDMY as compared to PC2 and PC3. Therefore, PC1 has the most dominant yield determinant having the positive loading coefficients for all the predictor variables.

### Soil Chemical properties

The improving soil fertility is critical for enhancing the maize yield and NUE and N fertilization into different splits along with biochar might be beneficial soil amendment for higher side of grain yield and NUE. SOC status changed with different splits of N and biochar treated plots over that of the initial status on surface soil (0–15 cm soil depth). The higher concentration of SOC with the N fertilizer (120 kg Nha^-1^) into 3 splits along with biochar (10 t ha^−1^) treated plots was a results of increased the higher root biomass and plant residues [38]. Additions of biochar along with N also increase microbial activity in the soil which may increase the soil organic carbon content in soil [63]. Hence it is proved that application of stable organic matter like biochar along the mineral fertilizer, not only increases soil fertility but also protects environment by its multifarious functions. Application of N rate (120 kg Nha^−1^) in 2 equal splits at V8 and VT growth stages also recorded higher values of NH_4_^+^-N (2.29 mg kg^−1^) and NO_3_^-^-N (0.44 mg kg^−1^) concentration as compared recommended dose of N and control treatment. This might be due to reduced N leaching losses which resulted higher N retention in the topsoil ultimately N recovery with the late split N application [45]. Furthermore, results showed that application of biochar (10 t ha^−1^) along with N fertilizer into multi-top dressed significantly increased the concentration of ammonium (NH_4_^+^-N) and nitrate (NO_3_^-^-N) in soil. The increase in NH_4_^+^-N and NO_3_^-^-N in the soil, this was due to high CEC of biochar and it potential to retain ammonium-N in the soil [64]. Since, biochar is more important as a soil conditioner and driver of nutrient transformations and it also minimized N leaching [65]. Similarly the concentration of NH_4_^+^-N and NO_3_^-^-N in the soil was increased as results of application of biochar along with N fertilizer might be due to decreasing N losses in the form of ammonia volatilization and denitrification and increasing N retention capacity of soils [66].

## Conclusion

Our three years research findings demonstrated significant improvement in maize yields and NUEs under the different agronomic management based on split nitrogen applications in rainfed maize in Vertisols of India. The recommended N management practices recorded lower yield, N uptake and NUEs compared to the agronomic based multi-split N application with skipping of basal N rates. The delayed N application at V8 and VT growth stages achieved higher grain yield, TDMY, N uptake, AE, PFP, PE and RE. Such delayed application has increased mean grain yield and TDMY by 13.5% and 11.2% over recommended practice, respectively. Pooled results illustrated that the late multi-split N application at V8 and VT recorded significant increases of 45.1%, 15.3%, 14.0% and 37.9% in AE, PFP, PE and RE than recommended practices, respectively. The grain yield, TDMY and NUEs were significantly increased with the application of biochar (10 t ha^−1^) along with recommended dose of N into 3 splits as compared to without biochar. Factor analysis showed positive and significant linear relationship between biomass, N uptake and biochar application which turned out to be more robust at V8 and VT growth stages of maize. Therefore, N uptake was greatest predictor of TDMY and AE was most strongly related to biomass production. Split application of N fertilizer significantly improved the SOC and NH_4_^+^-N and NO_3_^-^-N in the soil. Significant positive trend of SOC and available soil N status were observed under N fertilization (120kgNha ^−1^) along with biochar (10 t ha^−1^). Hence, the application of N rates into two equal splits at V8 and VT growth stages of maize with biochar application would be remunerative as alternative agronomic based approaches to synchronize the crop N demand and soil supply for higher crop yield and NUEs than the recommended practices in rainfed maize in Vertisols of India.

## Abbreviations

AE: agronomic efficiency;
DMA: dry matter accumulation;
N: nitrogen;
NDM: nitrogen content in dry matter;
NUDM: nitrogen uptake in dry matter;
NUE: nitrogen use efficiency;
PCR: principal Component Regression;
PE: physiology efficiency;
PFP: partial factor productivity;
RE: recovery efficiency;
STCR: soil test crop response;
TDMY: total dry matter yield;
VT: vegetative tasseling stage of maize.

## Acknowledgement

Authors are thankful to ICAR-Indian Institute of Soil Science (ICAR-IISS), Bhopal, for providing all assistance through an Institute project. We are grateful to Mr. Deepak Kaul, Chief Technical Officer and Mr. Jai Singh, Senior Technical Officer of the ICAR-IISS, Bhopal for their technical assistance provided with the field management and lab analyses conducted during the study period.

